# Genome-wide interrogation of genetic requirements for root colonization by CRISPRi-seq in *Bacillus subtilis*

**DOI:** 10.64898/2026.09.10.750580

**Authors:** Vikrant Minhas, Wiem Abidi, Jesús Cámara-Almirón, Huei-Hsuan Merissa Tsai, Valérie Dénervaud Tendon, Dimitra Synefiaridou, Jan-Willem Veening, Niko Geldner

## Abstract

*Bacillus subtilis* is a known plant growth promoting rhizobacterium. Yet, the genetic basis for effective *B. subtilis* root colonization remains unclear. Here, we applied genome-wide CRISPR interference sequencing (CRISPRi-seq) to systematically identify bacterial genes required for early *Arabidopsis thaliana* root colonization by the undomesticated *B. subtilis* strain NCIB3610. Complementary bacterial RNA-seq experiments in the same conditions further revealed a distinct transcriptional state of root-associated cells. Together, we identified 249 *B. subtilis* genes impacting root colonization. We reveal a critical, biofilm-independent, role of cell envelope remodeling during early root association, specifically wall teichoic acid synthesis and D-alanylation. Moreover, we observed a metabolic shift toward TCA-cycle-driven aerobic growth and utilization of fructose, induction of secondary metabolite biosynthesis, as well as active repression of sporulation, competence and prophage programs. Our work establishes CRISPRi-seq as a powerful tool for dissecting bacterial-plant interactions relevant to sustainable agriculture.

## Introduction

The complex interplay between plants and their associated microbiota is critical for plant health, development, and resilience to environmental stressors (1). Among the diverse array of beneficial microorganisms, the Gram-positive bacterium *Bacillus subtilis* has emerged as a prominent plant growth-promoting rhizobacterium (PGPR). This bacterium is widely studied for its ability to enhance plant growth, induce systemic resistance in plants and confer protection against phytopathogens (2–4). *B. subtilis* is thought to promote plant growth through several mechanisms, including the solubilization of phosphorus, indirectly increasing nitrogen availability (5), and production of compounds that modulate plant growth and development (5–7) such as phytohormones, or volatile organic compounds (8). It also induces systemic resistance in plants, bolstering their defenses against a spectrum of pathogens (9). The colonization of *Arabidopsis thaliana* (Arabidopsis) by *B. subtilis* can serve as a model system for investigating the mechanistic basis of the multi-faceted *B. subtilis* effects on plant growth and protection. *B. subtilis* is known to colonize the rhizosphere and root surfaces of Arabidopsis, forming biofilms that are critical for sustained root association. Biofilm formation is initiated by motile cells reaching and attaching to the root surface, followed by differentiation into matrix-producing cells. Swarming motility and the balance between motile and matrix-producing subpopulations, controlled by regulators such as Spo0A, SinR/SinI and SlrR, are also central to successful root colonization (10, 11). In addition, secondary metabolites produced during biofilm formation, including the lipopeptide surfactin, contribute both to cell spreading and to plant-defence priming (12). Moreover, bacterial colonization is enabled by root exudates, which provide particular nutrient environments for bacterial growth (10, 13).

The targeted approaches above have revealed important aspects for *B. subtilis* root colonization. Yet, to truly leverage the potential of *B. subtilis* for decreasing reliance on pesticides and fertilizers (14), it is essential to obtain a comprehensive understanding of the genetic determinants that facilitate *B. subtilis* colonization of plant roots. Here, we report the application of CRISPRi-seq to provide an unbiased, genome-wide view of the genetic determinants involved in *B. subtilis* root colonization. Recent advancements in functional genomics have enabled the development of CRISPR interference (CRISPRi) as a powerful tool for precise, genome-wide inducible gene repression in bacteria, allowing for the determination of essential genes in tested conditions (15). This approach utilizes a catalytically inactive Cas9 (dCas9) protein guided by a single-guide RNA (sgRNA) to obstruct transcription initiation, thereby facilitating systematic analysis of gene function at genome-wide scales (16). In contrast to transposon-based genetic screens such as Tn-Seq, CRISPRi is inducible, so knockdown can be timed and titrated rather than imposed at the moment of library construction. Essential genes, which are inaccessible to transposon mutagenesis by its design (17), therefore remain available to genome-wide screening, an advantage that matters wherever the processes under study depends on core cellular machinery. Genome-wide CRISPRi-seq has been implemented in a number of different bacterial species including *Escherichia coli, Staphylococcus aureus, Streptococcus pneumoniae, Mycobacterium tuberculosis, Haemophilus influenzae* and *B. subtilis*, among others (18–24). However, no genome wide CRISPRi-seq studies have been performed so far looking at bacterial interactions with the plant host. While Tn-seq has been performed to study Pseudomonadota-root interactions (25), a genome-wide screen (Tn-seq or CRISPRi) has not yet been performed in *B. subtilis* to the best of our knowledge. Here, we constructed a CRISPRi-seq library, targeting every gene and operon in the genome of the plant colonizing *B. subtilis* NCIB3610 strain, which effectively forms biofilms and colonizes plants (10). This library was applied to Arabidopsis to identify genes essential for root colonization. Our screen reveals several novel pathways important for root colonization, such as adaptation to oxidative stress, specific cell wall remodeling, or ABC-type export systems. Moreover, it provides comprehensive support for the importance of pathways previously implicated through single candidate gene knock-outs. Our screen is complemented with a RNA-seq profiling of root colonizing bacteria under the same conditions, further revealing surprising, specific carbon source utilization and cell wall remodeling responses, providing new insight into the molecular mechanisms underlying *B. subtilis*-plant interactions.

## Results

### *Construction of the* B. subtilis NCIB*3610 CRISPRi library*

A CRISPRi-seq library was created in the largely undomesticated *B. subtilis* strain NCIB3610 by inserting an IPTG-inducible *thrC::Pspank-dCas9-ery* cassette together with a constitutively expressed sgRNA library (*amyE* locus) targeting every gene and operon in the *B. subtilis* 168 genome (Figure 1A, Figure S1). The NCIB3610 strain is well studied for its ability colonize roots and form biofilms, making it an ideal candidate for this study (10, 26). Because *B. subtilis* 168 is >99% genetically identical to NCIB3610 across shared chromosomal regions, the library covers nearly every chromosomal gene. The notable exception is the ∼102 genes on the 3610-specific 84-kb pBS32 plasmid (absent from 168), which are therefore not represented (27). The library was tested for activation in vitro in LB +/- 1mM IPTG for ∼12 generations, with a substantial drop in sgRNA reads confirming successful induction (Figure S2A-C) and many known *B. subtilis* essential genes scoring as essential in the screen (Data S1, Figure S2D,E). As a targeted proof of concept, individual sgRNAs against two essential genes (*murF*, *dnaA*) and one non-essential gene (*ganA*) were tested in ½ MS liquid + 0.5% sucrose culture. Here, induction caused the expected growth defect only for the essential targets (Figure S2F), confirming functional knockdown in the 3610 background.

**Figure 1.**
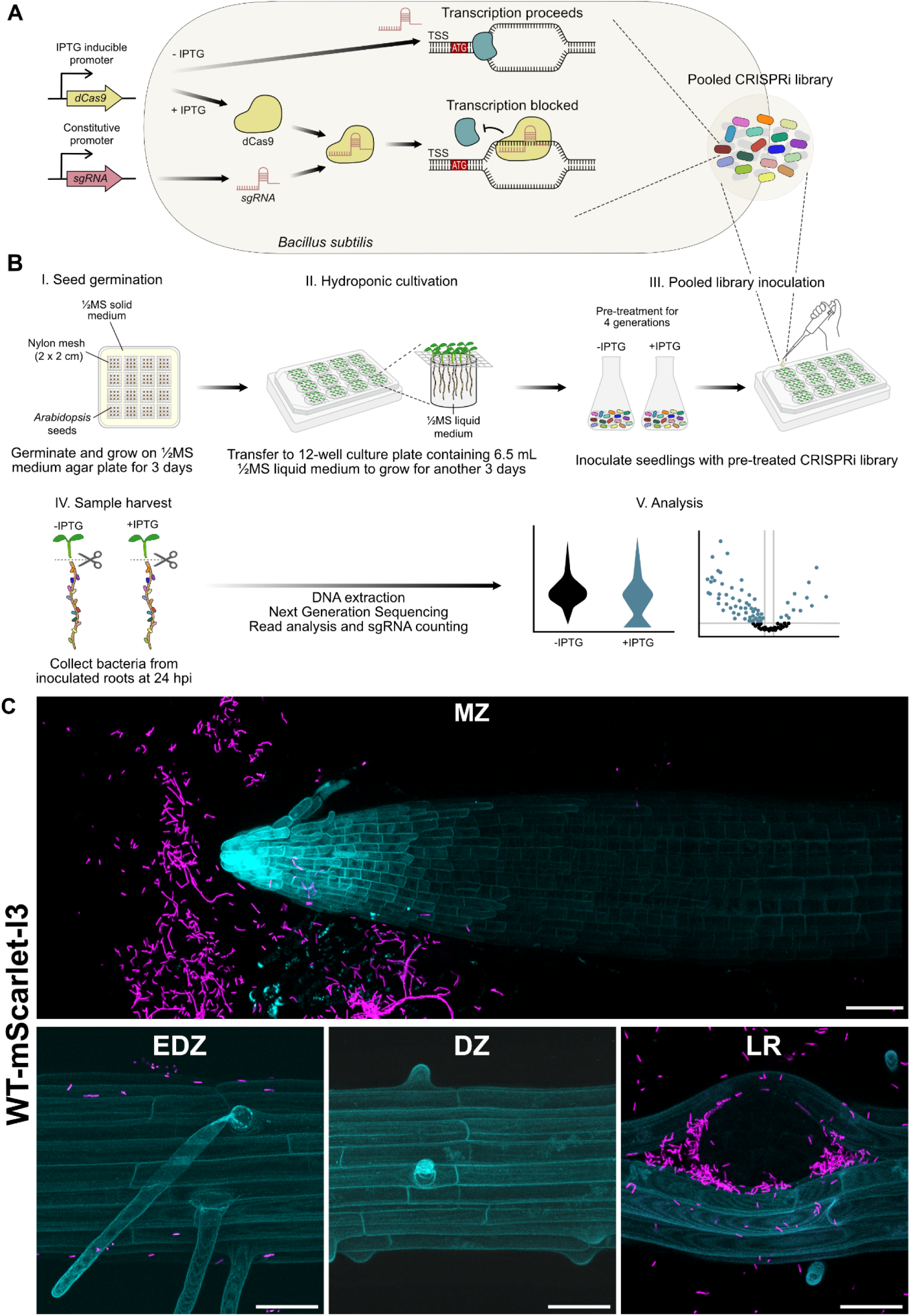
Schematic of *B. subtilis* CRISPRi-seq library construction and plant colonization workflow. (A) CRISPRi mechanism and design strategy. **(A)** An IPTG-inducible dCas9 and a constitutively expressed sgRNA targeting the region downstream of the transcriptional start site (TSS) are used to block transcription elongation. **(B)** Arabidopsis root colonization assay and CRISPRi-seq workflow. (I-II) Arabidopsis seeds are germinated on solid ½ MS medium for 3 days before transfer to a ½ MS hydroponic system, containing 1mM sucrose and 0.05mg/ml threonine, for an additional 3 days. (III) Seedlings are inoculated with the *B. subtilis* CRISPRi library (pre-treated with +/- IPTG). (IV-V) Bacteria are harvested from the roots and the liquid at 24 hours post-inoculation for DNA extraction and sgRNAs sequencing. Comparison of sgRNA abundance between conditions allows for the identification of genes essential for root colonization. **(C)** Confocal microscopy of root colonization with mScarlet-I3-tagged *B. subtilis* 3610 (magenta) on 6-day-old Arabidopsis seedling roots constitutively-expressing a mTagBFP2 protein in plasma membrane (cyan). Images are maximum-intensity z-projections, and are representative of n = 5 roots per strain from three independent experiments. MZ = meristematic zone, EDZ = early differentiated zone, DZ = differentiated zone, LR = lateral root. Scale bars: 106 µm for the MZ, 40 µm for all other zones.

Moreover, to assess library integrity and leakiness, sgRNA counts were examined in the uninduced condition in LB medium. Of the 4,326 sgRNAs, only 15 (0.35%) had zero read counts in at least one uninduced replicate, and 10 were absent from all four. The majority of these 15 missing sgRNAs target core components of the translation machinery, including nine ribosomal protein genes, and other central processes (Data S1). The same sgRNAs were already absent from the pre-induction libraries (13 zero-count sgRNAs, 12 shared with the LB screen), placing this depletion during library propagation rather than during the screen itself. Together, this pattern is consistent with a very low leakiness of dCas9 expression during library propagation, which selectively depletes very few sgRNAs, that are targeting the most essential cellular processes (28).

### *CRISPRi-seq of* B. subtilis *during root colonization*

To utilize the 3610 CRISPRi-seq library to study colonization dynamics of *B. subtilis* on Arabidopsis roots, a mini-hydroponics system was set up for *Arabidopsis* seedlings (Figure 1B). To test this system, 6-day old Arabidopsis wild-type seedlings, containing a constitutive Lti6b-mTagBFP2 plasma membrane marker (29), were inoculated with a consitutively expressed mScarlet-I3-fluorescent 3610 strain at OD_600_ 0.02 for 24h in ½ MS plant growth medium + 1mM sucrose + 0.05mg/ml of threonine, before roots were imaged with confocal fluorescence microscopy (see materials and methods). Here, *B. subtilis* showed clear spatial discrimination, as it colonized root cap cells and emerging lateral roots in high numbers, while they were present in only low amounts throughout the rest of the root (Figure 1C). It is known that for successful *B. subtilis* colonization of Arabidopsis roots in hydroponics, small concentrations of a carbon source are required (30). We found that glycerol, the carbon source traditionally used in Arabidopsis - *B. subtilis* interaction studies (10, 31), caused significant root stunting in our ½ MS system, consistent with its reported inhibitory effects on Arabidopsis root development (32). Sucrose, by contrast, did not deleteriously affect root development (Figure S3, Data S2). Hence, sucrose was used as a more plant-compatible carbon source in all subsequent experiments.

The 3610 CRISPRi-seq library was pre-induced with 1 mM IPTG for ∼4 generations in LB before inoculation at OD_600_ 0.02 with Arabidopsis seedlings in the hydroponic system, with 2 mM IPTG added at 0 h and 6 h post-inoculation to ensure library induction. Pre-induction did not significantly drop sgRNA representation (Figure S4, Data S3), but was necessary for successful library induction on roots, displaying substantial drops in sgRNA reads and good variation between conditions (Figure S5). To identify genes required specifically for root colonization, root-attached and free-swimming bacteria in the liquid medium were collected 24 h post-inoculation and sgRNA reads compared between conditions (Figure 2A-C, Data S4). 235 genes were significantly required (log_2_FC < -1, padj < 0.05) for root colonization, while 14 sgRNAs were significantly costly (log_2_FC ≥ 1, padj < 0.05). Numbers in parentheses after gene names in the following results indicate log_2_FC. Gene set enrichment analysis (GSEA) on SubtiWiki categories (33) identified 12 categories significantly enriched for root colonization (Data S5), including flagellar proteins, swarming, and biosynthesis of teichoic acids and isoprenoids (Figure 2D).

**Figure 2.**
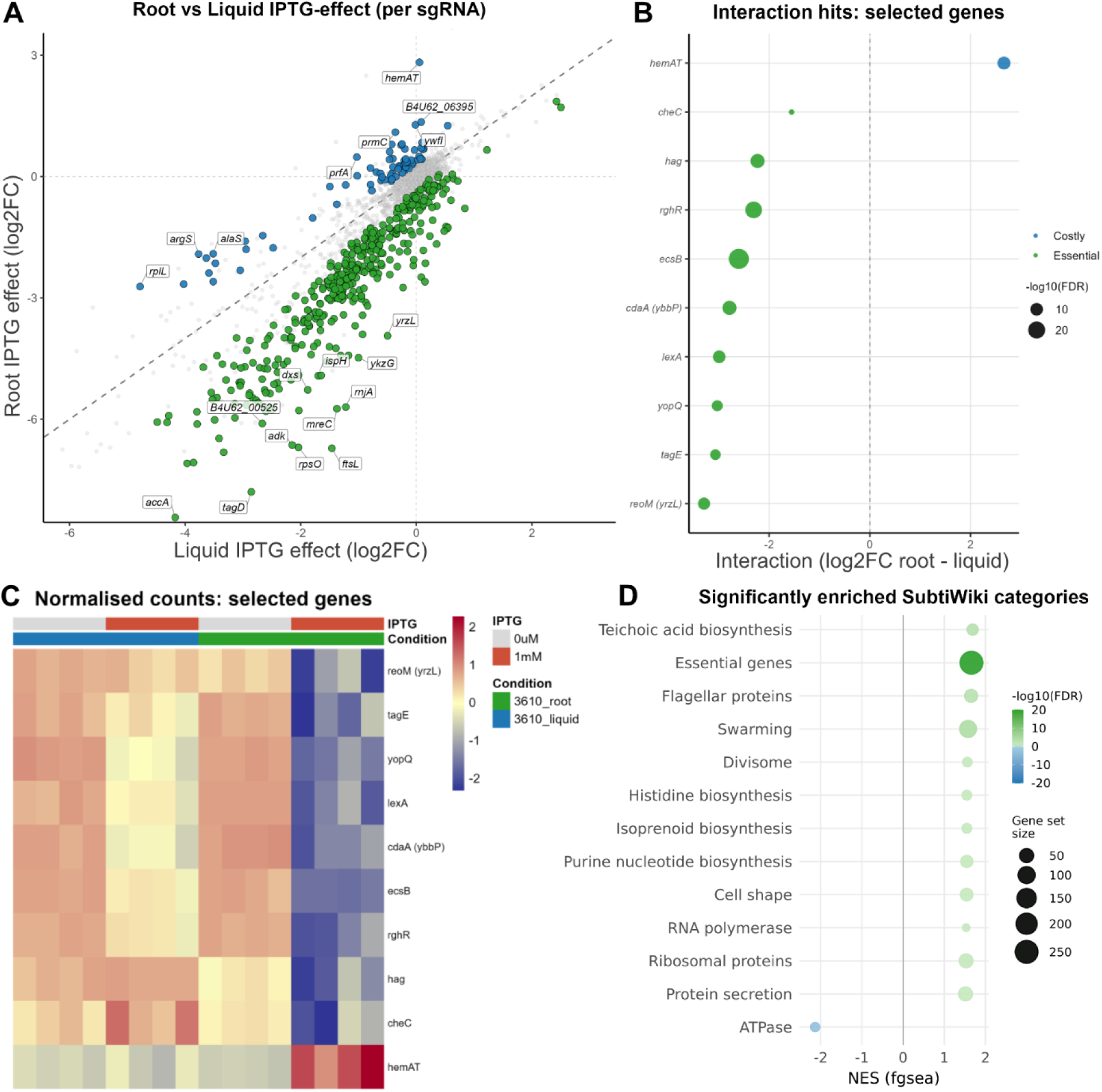
CRISPRi-seq analysis of *B. subtilis* colonization on *Arabidopsis* roots. **A)** Scatter plot comparing the IPTG-induction effect on sgRNA fitness in liquid (x-axis, log_2_FC 1 mM vs 0 µM) versus root-colonizing conditions (y-axis, log_2_FC 1 mM vs 0 µM). Each point represents one sgRNA. Green points indicate sgRNAs with a significantly stronger knockdown effect in root (root-essential; FDR < 0.05), blue points indicate sgRNAs with a significantly stronger knockdown effect in liquid (root-costly; FDR < 0.05), and grey points are non-significant. The dashed diago-nal represents the identity line (x = y). Labels show the top hits by interaction effect size. **B)** Ranked dot plot showing selected interaction hits by interaction log_2_FC (root - liquid). Each dot represents one gene; colour indicates direction (green = essential, blue = costly in root) and dot size represents -log10(FDR). **C)** Heatmap of DESeq2 normalized counts (log2-transformed and z-scored per gene) for the selected interaction hit genes. Columns represent individual biological replicates ordered by condition (liquid 0 µM → liquid 1 mM → root 0 µM → root 1 mM), with four biological replicates per condition; each replicate comprises 108 pooled seedlings. Annotation bars indicate IPTG concentration and growth condition. **D)** Bar chart showing significantly enriched SubtiWiki functional categories (FDR < 0.05) from gene set enrichment analysis (GSEA) of the interaction term. Bar colour indicates direction (green = enriched in root-essential genes, blue = enriched in root-costly genes) and colour intensity encodes -log10 (FDR). The x-axis represents the normalised enrichment score (NES).

Inspection of individual sgRNA hits provided further resolution within these categories. The 235 required genes mapped onto four broad functional groups: motility and chemotaxis, cell envelope biogenesis and cell division, the isoprenoid/MEP pathway and a smaller set of regulatory and stress-response loci. Among the strongest fitness defects were core flagellar/chemotaxis components, including *hag* (−2.23), *fliD* (−2.67), *sigD* (−1.34), *cheC* (−1.55) and *yopQ* (−3.03). The cell-envelope category was particularly densely populated and tightly clustered: wall teichoic acid (WTA) biosynthesis genes *tagA* (−3.03), *tagD* (−5.28), *tagE* (−3.06) and *tagF* (−1.58) were all significantly depleted, alongside the peptidoglycan and cell-shape determinants *mreC* (−3.42), *pbpB* (−2.41) and *reoM*/*yrzL* (−3.29), and the cell-division components *ftsL* (−5.62), *divIC* (−2.80) and *minC* (−1.15). Consistent with the requirement for undecaprenyl-phosphate as a WTA carrier, the MEP isoprenoid pathway required for its synthesis was co-essential, with *dxs* (−3.10), *ispD* (−1.78) and *ispH* (−3.13) all depleted. Beyond these categories, several individual loci stood out: the c-di-AMP synthetase *cdaA*/*ybbP* (−2.78), the SOS repressor *lexA* (−2.99), the sporulation/biofilm regulator *rghRA* (−2.31) and the ABC transporter *ecsB* (−2.60). On the costly side, only 14 sgRNAs were significantly enriched on roots, and these clustered into two interpretable groups. The first centered on aerotaxis and heme metabolism: the soluble oxygen-sensing chemoreceptor *hemAT* (+2.66) was the single strongest costly hit, accompanied by the heme biosynthetic gene *ywfI* (+1.19) and the mechanosensitive channel *mscL* (+1.15). A second, less expected cluster concerned translation: four aminoacyl-tRNA synthetases were costly when knocked down (*argS* +1.71, *alaS* +1.49, *valS* +1.07, *gltX* +1.15,), together with release factor *prfA* (+1.27) and the ribosomal protein *rplL* (+1.46). We also corrected our CRISPRi-seq calls for the different numbers of generations that root-attached and free-swimming populations may complete. This more stringent comparison reduced the sgRNAs called required from 235 to 80, with the loss concentrated in growth-essential genes, but left motility as the dominant signature (swarming, FDR = 9.3 × 10^-4^; flagellar proteins, FDR = 9.2 × 10^-5^) (Figure S6, Data S7). Importantly, every CRISPRi-seq hit chosen for validation below remained significant.

### Validation of CRISPRi-seq hits

Several CRISPRi-seq hits with known or potential roles in root colonization were selected for individual validation (Figure 2B,C): cell motility (*hag*, *yopQ*) (34, 35), chemotaxis (*cheC*) (36, 37), DNA damage response (*lexA*) (38), cyclic di-AMP synthesis (*ybbP*/*cdaA*) (39), cell wall synthesis (*tagE*, *yrzL*/*reoM*) (40–42), as well as the ABC transporter *ecsB*, involved in cell division and stress adaptation, and *rghR*, a modulator of quorum sensing and surfactin production (43, 44). We additionally selected *hemAT* (45), the soluble oxygen-sensing chemoreceptor that scored as the strongest costly hit. Each gene was deleted in *B. subtilis* 3610 by homologous recombination of a kanamycin-resistance cassette. These mutant strains and the WT were then inoculated onto *Arabidopsis* roots for 24h, both individually and in competition with the WT, in the same manner as in the CRISPRi-seq experiment, before bacterial colonization on roots was enumerated via colony counting. All of the mutant strains of the CRISPRi-seq hits were affected in their ability to colonize roots when compared to 3610 WT (Data S6), both individually (Figure 3A) and in competition with WT 3610 (Figure 3B). Interestingly, Δ*hemAT* showed a significant colonization defect when inoculated individually (Figure 3A). However, when co-inoculated in direct competition with WT 3610, the mutant showed significantly higher CFUs relative to WT (Figure 3B), suggesting that the Δ*hemAT* advantage is contingent on the presence of WT cells

Since we had observed spatial preference of bacterial colonization at the root surface, we did imaging of mutant strain colonization in competition with wild-type (Figure 3C). For both *reoM* and *tagE*, we found the same spatial patterns of colonization and confirmed clearly reduced numbers GFP-labelled, mutant bacteria. By contrast, we observed that, in competition with wild-type, Δ*hemAT* does significantly better at the root cap than at the lateral root emergence site, suggesting a spatial niche-specific difference in Δ*hemAT* colonization that cannot be observed by CFU counting assays. This preference could indicate different oxygen levels between these two niches, for example. However, our image-based quantification did not show the expected, overall enhanced colonization capacity of Δ*hemAT* compared to wild-type, with pixel ratios of Δ*hemAT/WT* below one for both niches (Figure 3D, Data S8). Given that both CRISPRi-seq and CFU count data of *hemAT* knock-outs both outcompeting WT, this discrepancy may be explained by a difference in cell size and motility that we observe between mutant and wild-type colonies, with wild-type forming a markedly higher proportion of non-motile, chained and elongated cells. Because our imaging-based quantification relies on fluorescent pixel counting rather than direct cell counts, such chaining would amplify the apparent wild-type signal relative to the smaller, more dispersed and more motile mutant cells.

**Figure 3.**
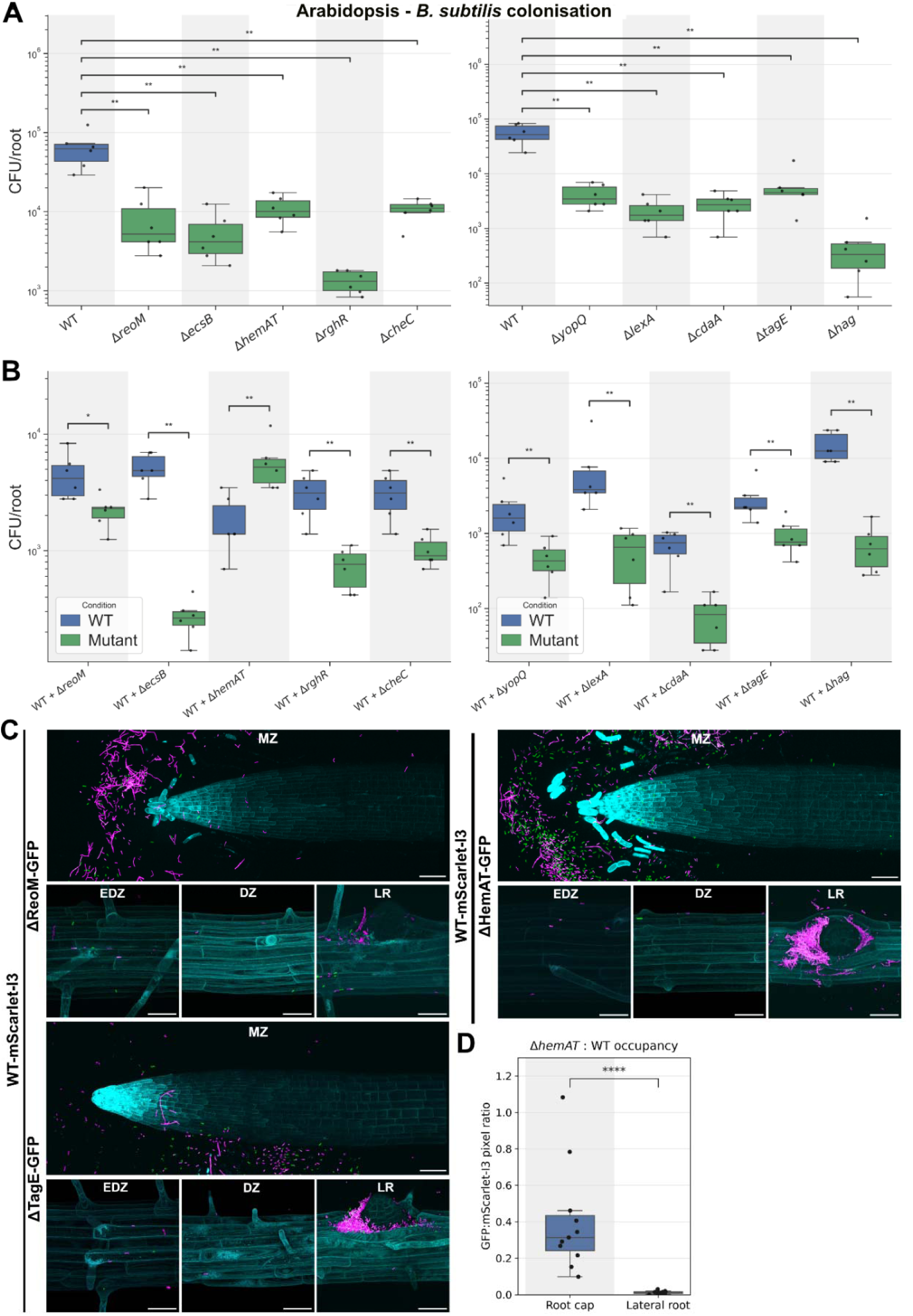
Validation of *B. subtilis* root colonization genes and phenotypes. **A)** Individual root colonization assay. CFU counts per root of WT 3610 and knockout mutants recovered from Arabidopsis roots 24 h after individual inoculation at OD_600_ 0.02 in the hydroponic system. **B)** Competitive root colonization assay. CFU counts per root of each mutant (green) and WT (blue) recovered from the same root homogenate 24 h after co-inoculation at equal starting amounts (OD_600_ 0.01 per strain), enumerated by selective plating. In A and B, boxes show the median and interquartile range; n = 6 biological replicates per strain, each of nine pooled roots, from two independent experiments. Comparisons (A, each mutant versus WT; B, mutant versus WT within each co-culture) were by exact two-sided Mann-Whitney U test with Benjamini-Hochberg adjustment within each panel (* adjusted p < 0.05, ** adjusted p < 0.01). **C)** Confocal microscopy of competitive root colonization. Wild-type *B. subtilis* tagged with mScarlet-I3 (magenta) was co-inoculated with GFP-tagged Δ*reoM*, Δ*hemAT* or Δ*tagE* (green) at equal starting amounts (OD_600_ 0.01 per strain) and imaged 24 h post-inoculation on 6-day-old Arabidopsis seedling roots constitutively-expressing a mTagBFP2 protein in plasma membrane (cyan). Images are maximum-intensity z-projections, spectrally unmixed to separate GFP and mScarlet-I3 emission, and are representative of n = 5 roots per strain from three independent experiments. MZ = meristematic zone, EDZ = early differentiated zone, DZ = differentiated zone, LR = lateral root. Scale bars: 106 µm for the meristematic zone, 40 µm for all other zones. **D)** Relative abundance of Δ*hemAT* at the root cap (blue) and lateral root emergence sites (green) in competition with WT, expressed as the ratio of GFP (Δ*hemAT*) to mScarlet-I3 (WT) above-threshold pixel counts per confocal image, from roots co-inoculated and imaged as in C. Boxes show the median and interquartile range; points show individual roots. n = 11 roots per niche, from three independent experiments. Niches were compared by exact two-sided Mann-Whitney U test (**** p < 0.0001).

### RNAseq of B. subtilis during root colonization

To complement the CRISPRi-seq screen and capture the transcriptional dimension of root colonization, we performed RNA-seq on *B. subtilis* recovered from the same hydroponic system. Bacteria were collected 24 h after inoculation, with root-attached cells compared directly against the liquid free-swimming population in the surrounding medium (Figure S7). Differential expression analysis identified 400 genes significantly upregulated and 833 genes significantly downregulated on roots (Figure 4A,B) (Data S9), corresponding to roughly a quarter of the genome and indicating a pronounced re-shaping of the transcriptional state on contact with Arabidopsis. The differentially expressed genes resolved into six biological themes: (1) a metabolic shift toward TCA-cycle-driven aerobic growth fed by fructose and oligopeptide uptake, (2) a coordinated cell envelope remodeling including teichoic-acid D-alanylation, (3) an active suppression of sporulation, competence and prophage pathways, (4) an induction of secondary metabolite and biofilm matrix production, (5) an upregulation of stress and detoxification responses and (6) a repression of siderophore-based iron scavenging.

**Figure 4.**
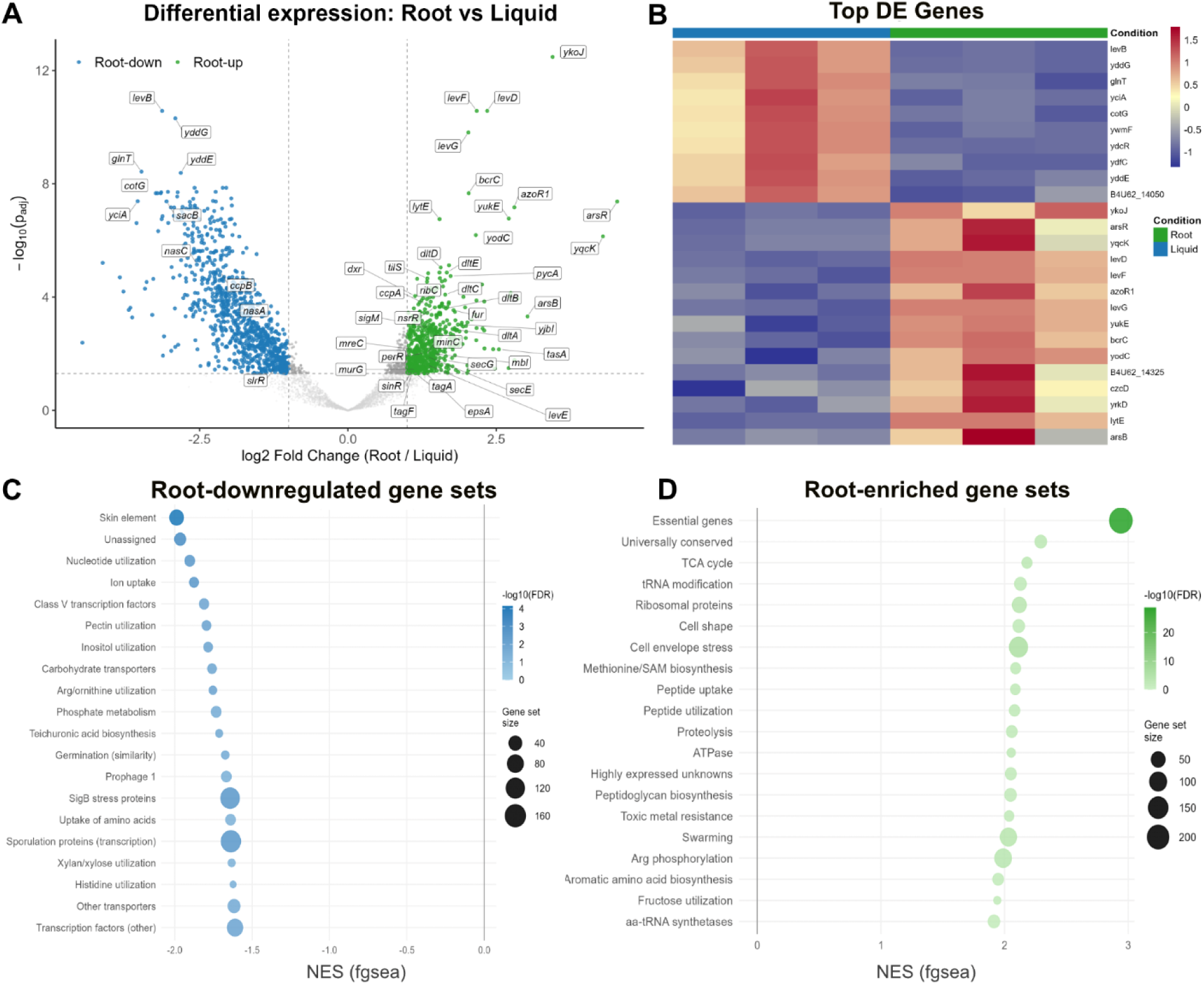
Transcriptomic analysis of *B. subtilis* during Arabidopsis root colonization. Wild-type *B. subtilis* 3610 was inoculated into the Arabidopsis hydroponics system. After 24 hours, RNA was isolated from root-colonizing and liquid non-colonizing bacteria and analysed by RNA-seq using DESeq2. **A)** Volcano plot showing differential gene expression between root-colonizing and liquid non-colonizing *B. subtilis*. Green dots represent genes significantly upregulated in root, blue dots represent genes significantly upregulated in liquid (|log₂FC| ≥ 1, adjusted p-value < 0.05). Grey dots are non-significant. Dashed lines indicate fold-change and significance thresholds. Labels show key genes of interest; black italic text with white background boxes highlights named genes. **B)** Heatmap of regularized-log (rlog) transformed counts (z-scored per gene) for the top differentially expressed genes (top 15 root-upregulated and top 10 root-downregulated by log_2_FC × -log_₁₀_(padj)). Columns represent individual biological replicates ordered by condition (liquid → root), with three biological replicates per condition retained for analysis. The annotation bar indicates condition (green = root, blue = liquid). **C)** Dot plot showing significantly enriched SubtiWiki functional categories in root-colonizing *B. subtilis* (NES > 0, FDR < 0.05). Each dot represents one gene set; dot size encodes gene set size and colour intensity encodes -log10(FDR) on a green gradient. The x-axis represents the normalised enrichment score (NES). **D)** Dot plot showing significantly enriched SubtiWiki functional categories in liquid non-colonizing *B. subtilis* (NES < 0, FDR < 0.05). Dot size encodes gene set size and colour intensity encodes -log10(FDR) on a blue gradient. The x-axis represents the normalised enrichment score (NES).

On roots, *B. subtilis* switched into an apparent fructose-fueled aerobic growth state. This is indicated by the fructose phosphotransferase operon *levDEFG* being induced (*levD* +2.40, *levE* +1.31, *levF* +2.08, *levG* +2.03), together with the fructosamine deglycase *frlB* (+2.22) and the catabolite-control regulator *ccpA* (+1.34) (Figure 4A,B). In parallel, the endolevanase *levB* (− 3.08) was strongly downregulated and pathways for complex plant polysaccharide breakdown were coordinately repressed, with “utilization of pectin” among the significantly enriched root-downregulated categories in the GSEA (Figure 4C, Figure S8). Together with the downregulation of *sacB*, this points to monomeric fructose rather than sucrose or polymeric fructans as the primary carbon source (46–48). The full TCA cycle was induced, ATP synthase subunits and high-affinity aerobic cytochrome oxidases were upregulated, while the microaerobic cytochrome *bd* oxidase (*cydABCD*, all -1.70 to -2.00) and the assimilatory nitrate reductase (*nasABC*, all -1.66 to -2.82) were strongly repressed. Nitrogen acquisition shifted in parallel toward exogenous peptides as the oligopeptide and dipeptide transporter operons (*appBCDF*, *dppA-E*) were broadly induced (+1.33 to +1.67).

Cell-envelope adaptation was equally prominent at the transcriptional level. The entire *dlt* D-alanylation operon was induced (*dltA-E*, all +1.37 to +1.66), alongside the cell wall and division gene *mreC* (+1.04), fitting the requirement for cell envelope genes seen in the CRISPRi-seq screen. Markers of nitrosative stress also appeared, with the truncated haemoglobin *yjbI* (+1.85) and the NO-responsive repressor *nsrR* (+1.44) significantly upregulated. Concurrently, the sporulation, germination, competence and prophage regulons (including SPβ and PBSX) were strongly repressed, indicating that root-attached cells were committed to vegetative growth, in contrast to planktonic cells entering stationary-phase programmes. A second clear signal came from secondary-metabolite and biofilm-matrix loci. The surfactin, plipastatin and bacillaene biosynthetic operons were coherently upregulated on roots (*srfAA* +1.61, *srfAC* +1.90, *srfAD* +1.80; *ppsE* +1.62; *pksS* +1.55), as was the matrix protein-fiber gene *tasA* (+2.59), consistent with the established role of these molecules in root association. Unexpectedly, the iron-scavenging machinery moved in the opposite direction: siderophore biosynthesis (*dhbE* -1.65) and uptake (*feuA* -1.50, *feuB* -1.37, *feuC* -1.64; *fhuB* – 1.18*, fhuG* -1.10) were among the most strongly repressed gene sets, suggesting that iron is not rate-limiting in the rhizosphere of hydroponically grown roots. A GSEA on the RNA-seq dataset confirmed these patterns at the pathway level, with two of the highest normalized enrichment scoring categories being Cell shape and Cell envelope stress proteins, paralleling the Biosynthesis of teichoic acids category from the CRISPRi-seq dataset, and Swarming, which also enriched in the CRISPRi-seq GSEA (Figure 4D, Data S10).

The largest transcriptional response on roots was a metal detoxification programme. The arsenic-responsive regulator *arsR* (+4.50) was the most strongly induced gene in the entire dataset, accompanied by the ArsR regulon members *arsB* (+3.02) and *czcD* (+2.78), and by copper homeostasis genes *copA* (+1.72) and *copZ* (+1.52) (Figure 4B, Data S7). The cannibalism effectors, *skfA* (+2.52) and *sdpC* (+2.21), were also induced, although the structural and immunity components of their respective operons were not. Nitrogen physiology adjusted in a similar direction with the sodium-glutamine symporter *glnT* being the sixth most strongly downregulated gene overall (−4.15), and other amino-acid catabolism genes followed, including *hutH* (−2.67) and *rocG* (−2.89), alongside the already mentioned repression of nitrate assimilation (*nasABC*). This pattern is consistent with nitrogen being acquired predominantly via the induced oligopeptide transporters rather than from inorganic sources or amino-acid catabolism. Two cofactor biosynthesis programmes were also induced, with the entire biotin operon upregulated (*bioWAFDBI*, all +1.32 to +1.68), and the methionine salvage pathway was partially induced (*mtnK*/*A*/*D*/*W*/*X*/*U*/*B*, all +0.90 to +1.81). This likely reflects genuine cofactor limitation at the root surface as neither biotin nor the methionine-cycle intermediates are present in ½ MS medium, and high demand is expected given the essentiality of the biotin-dependent carboxylases *accABC* and *pycA* in the CRISPRi-seq screen.

### CRISPRi-seq vs RNA-seq data

Comparing and integrating the CRISPRi-seq and RNA-seq datasets (data S11) provides a synoptic view of the functional and transcriptional landscape of root colonization. Gene set enrichment analysis revealed a discrete cluster of categories that were significant in both assays (Figure 5A). These were Biosynthesis of teichoic acids, Cell shape, Ribosomal proteins, DNA replication, Swarming, and Essential genes, showing simultaneous induction on roots and requirement for fitness, therefore defining a high-confidence core of pathways required for competitive root colonization.

**Figure 5.**
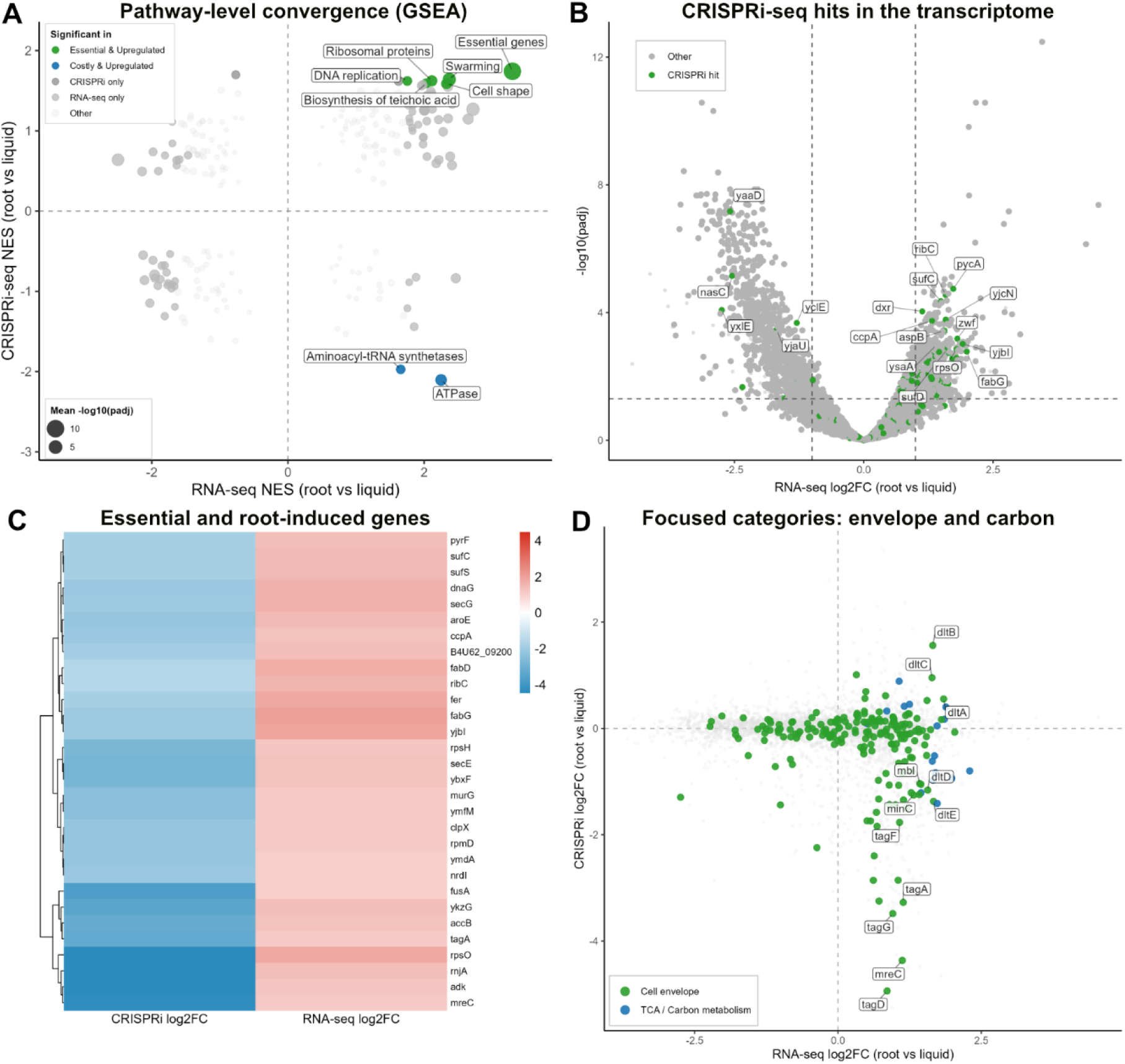
Comparative analysis of essentiality and transcriptomics during Bacillus subtilis root colonization (CRISPRi-seq vs. RNA-seq). Both screens compare root-colonizing to liquid, non-colonizing conditions. **A)** Pathway-level convergence between CRISPRi-seq fitness and RNA-seq expression. GSEA was performed independently on both datasets using SubtiWiki functional categories. Points show each category’s normalized enrichment score (NES) in RNA-seq (x-axis) vs. CRISPRi-seq (y-axis) GSEA; size reflects mean -log10(padj). Categories significant in both analyses (FDR < 0.05) are highlighted in blue or green and labeled; all others are grey. **B)** Volcano plot highlighting CRISPRi-seq hits within RNA-seq data. RNA-seq log_2_FC (root vs. liquid) vs. -log10(padj); genes meeting CRISPRi-seq hit criteria (FDR < 0.05, log_2_FC ≥ 1) are green, others grey. Dashed lines mark RNA-seq significance (padj = 0.05) and effect-size (log_2_FC = 1) thresholds. Labels show the top 18 genes significant in both screens, ranked by combined RNA-seq significance and effect size. **C)** Heatmap of genes with convergent essentiality and induction. log_2_FC (CRISPRi-seq and RNA-seq) for genes meeting significance thresholds (FDR < 0.05, log_2_FC ≥ 1) in both screens and showing the convergent direction: essential in CRISPRi-seq, upregulated in RNA-seq. Rows are hierarchically clustered by log_2_FC similarity. Blue = depletion/essentiality or downregulation; red = enrichment/dispensability or upregulation. **D)** Focused comparison of cell envelope and carbon metabolism genes. CRISPRi-seq log_2_FC (y-axis) vs. RNA-seq log_2_FC (x-axis), restricted to two functional categories: cell envelope biogenesis (green) and TCA cycle/carbon metabolism (blue); other genes shown in grey. Labels are restricted to genes within these categories meeting CRISPRi-seq or RNA-seq significance thresholds.

Nevertheless, overall genome-wide correlation between the RNAseq and CRISPRi datasets is weakly negative (r = -0.185, Figure S9A), and only 1.1% of all genes (n =46 were significant in both screens simultaneously (Figure S9C), suggesting that dual significance in both assays is highly stringent and selective. Indeed, many constitutively-expressed genes can nevertheless be required for root colonization. Moreover, despite being harvested at the same time point, the two datasets capture fundamentally different temporal windows: CRISPRi-seq integrates cumulative competitive colonization success across the entire 24 hour experiment, while RNA-seq provides a snapshot of the transcriptional state only at the endpoint. A gene can therefore be transcriptionally induced at 24 hours without having contributed to fitness over the colonization period. Conversely, a gene can be essential for early colonization steps without being transcriptionally active by the time of harvest. The motility-to-sessility transition is an illustration of this: flagellar genes are among the most essential in CRISPRi-seq, motility is required to initially reach the root surface, yet are strongly repressed in RNA-seq at 24 hours (Figure 5A, Figure S9), consistent with the well-described suppression of flagellar gene expression in *B. subtilis,* once they have reached the surface.

Adaptation of the cell envelope appears as a dominant process for root colonization, in both datasets. The RNA-seq volcano (Figure 5B) shows that CRISPRi hits (green dots) are disproportionately enriched among genes induced on roots, particularly in the cell envelope, stress response and secondary metabolite clusters. Figure 5C shows the top 30 genes that are simultaneously fitness-depleted in CRISPRi-seq (blue, left column) and transcriptionally-induced in RNA-seq (red, right column). Cell envelope genes (*tagA*, *mreC*, *murG*, *secE*, *secG*), ribosomal proteins (*rpsH*, *rpsO*, *rpmD*), translation factors (*fusA*) and metabolic genes (*accB*, *adk*, *fabD*, *fabG*) dominate this concordant set. This pattern is revealed at the gene-set level in Figure 5D, which colours all genes for the two most outstanding functional categories: cell envelope genes (green) cluster tightly in the essential-and-induced quadrant (negative CRISPRi log_2_FC, positive RNA-seq log_2_FC), while TCA/carbon metabolism genes (blue) are more broadly distributed. WTA biosynthesis genes *tagA* and *tagF*, peptidoglycan and cell-shape determinants *murG*, *mreC* and *mbl*, components of the general secretion machinery (*secE*, *secG*) and the cell-division gene *minC*, were both significantly upregulated in RNA-seq and significantly fitness-depleted in CRISPRi-seq. The bacterium therefore both scales up envelope synthesis transcriptionally and cannot tolerate its reduction functionally, identifying the cell envelope adaptation as a crucial step for early root colonization.

### Cell wall remodeling mutants

Both datasets converged on the teichoic acid pathway, but with different gene sets. Eight sgRNAs targeting wall teichoic acid biosynthesis (*tagA, tagB, tagD, tagE, tagF, tagG, tagH, tagO*) scored in the CRISPRi-seq screen. However, the *dlt* operon, which D-alanylates teichoic acids, was not identified in CRISPRi-seq, yet *dltA-E* were strongly induced on roots in the RNA-seq experiment (Figure 4). Since *dlt* sgRNAs are present at very low abundance in the CRISPRi-seq screen (baseMean 3.6 *dltB*, 10.2 *dltA*, 12.6 *dltD*, 23.2 *dltC*, 34.6 *dltE*), the screen was underpowered to detect their depletion, possibly explaining this discrepancy. Hence, we decided to generate a *dltA* knock-out mutants, likely affecting the entire *dlt* operon, and indeed found that *dltA* or the *dlt* operon is also necessary for root colonization in our assays, both in mono-association and in competition with wild-type (Figure S10A,B).

To test whether cell wall composition contributes to colonization at the earliest stage, before biofilm formation, we performed direct attachment assays. Six-day-old Col-0 seedlings were inoculated with wild type, Δ*tagE*, Δ*reoM* or Δ*dltA*, and adherent bacteria quantified after 30 min and 2 h. We found all three mutants to attach significantly less to roots than the WT both at 30 min and 2h post-inoculation (Figure 6A, data S12). D-alanylation is the principal means by which *B. subtilis* modulates the net negative charge of its teichoic acids, which might be necessary to attach to pectin-containing, negatively-charged plant cell walls. We therefore asked whether the Δ*dltA* defect could be reproduced on a chemically-defined, abiotic surface. Bacteria were presented with negatively charged polygalacturonic acid (PGA), the principal acidic pectin of the plant cell wall, adsorbed onto polystyrene wells. Indeed, Δ*dltA* attached to PGA significantly less than WT 3610 (Figure 6B, data S13), closely matching its defect on roots. On positively charged poly-L-lysine surfaces or uncoated polystyrene, Δ*dltA* did not show significant differences (Figure S10D,E). An inert polyanion therefore reiterates the root-attachment defect of Δ*dltA*, supporting a model whereby *B. subtilis* adapts its cell wall to optimize attachment to plant cell walls and assist in early root establishment.

**Figure 6.**
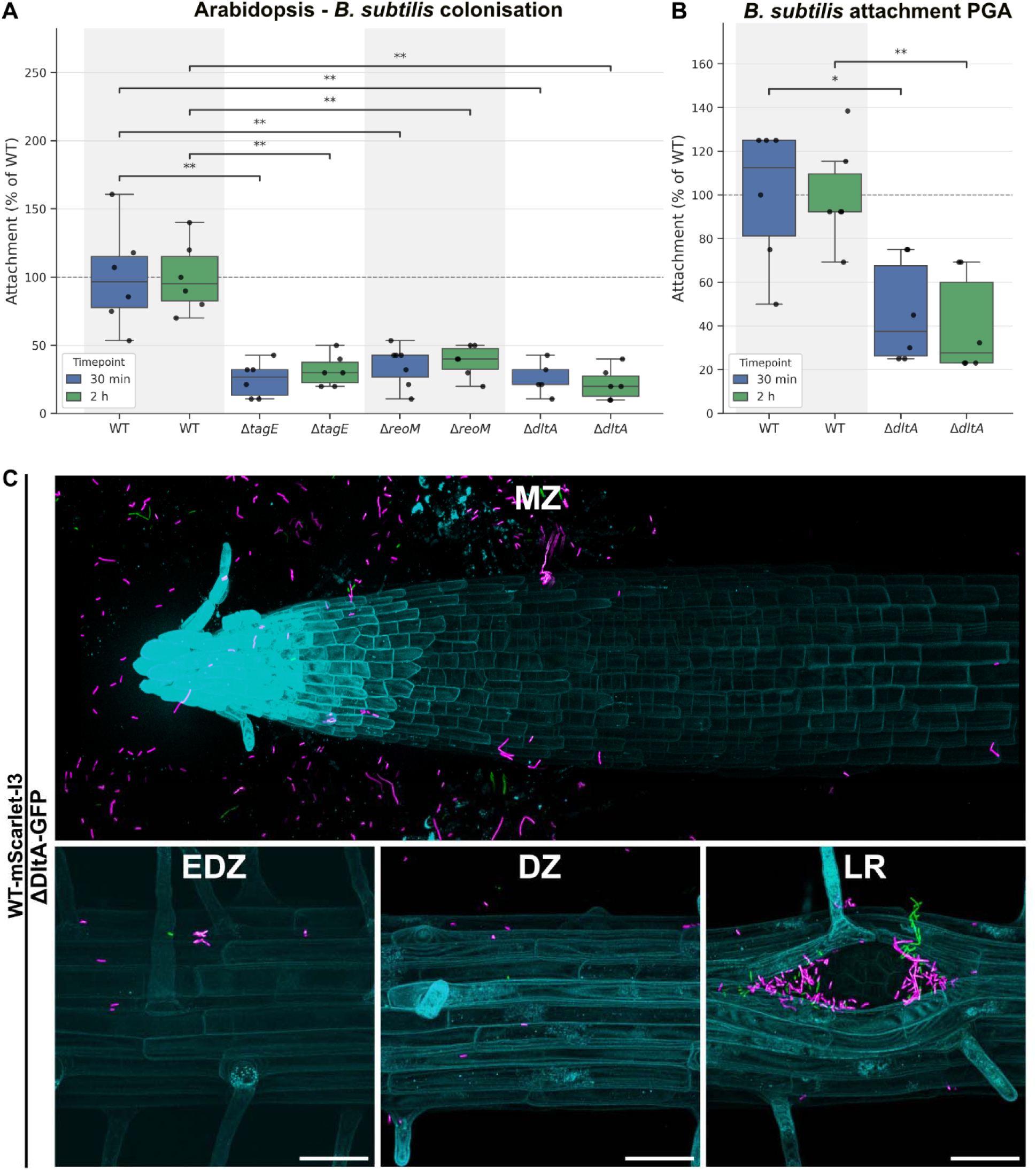
Early attachment to *Arabidopsis* roots and to a pectin-coated surface. **(A)** Six-day-old Col-0 seedlings grown vertically on ½ MS agar were inoculated with 10 µL of wild-type, Δ*tagE*, Δ*reoM* or Δ*dltA B. subtilis* at OD_600_ 0.02. Roots were washed and adherent bacteria enumerated by CFU counting after 30 min (blue) and 2 h (green), before biofilm formation. **(B)** Wild type and Δ*dltA* attached to polygalacturonic acid (PGA)-coated polystyrene wells in base assay buffer (10 mM sodium phosphate pH 7.0, 50 mM NaCl), assayed over the same two intervals. Values are expressed as a percentage of the mean wild-type value within each timepoint; the dashed line marks wild-type attachment. n = 6 biological replicates per group, from two independent experiments of three replicates each; one biological replicate comprises nine pooled seedlings in (A). Each mutant was compared with the wild type within its time point by exact two-sided Mann-Whitney *U* test, with *p*-values adjusted within the panel by the Benjamini-Hochberg procedure (* adjusted *p* < 0.05, ** adjusted *p* < 0.01; ns, not significant). **(C)** Confocal microscopy of competitive root colonization by ΔdltA. Wild-type *B. subtilis* tagged with mScarlet-I3 (magenta) was co-inoculated with GFP-tagged Δ*dltA* (green) at equal starting amounts (OD_600_ 0.01 per strain) and imaged 24h post-inoculation on 6-day-old Arabidopsis seedling constitutively-expressing a mTagBFP2 protein in plasma membrane (cyan). Images are maximum-intensity z-projections, spectrally unmixed to separate GFP and mScarlet-I3 emission (see Materials and Methods), and are representative of n = 5 roots from three independent experiments. MZ = meristematic zone, EDZ = early differentiated zone, DZ = differentiated zone, LR = lateral root. Scale bars: 106 µm for the meristematic zone, 40 µm for all other zones.

Electrostatic repulsion should be sensitive to ionic strength, since salt shortens the distance over which surface charges interact. Indeed, we found that the Δ*dltA* defect, significant at 0 and 50 mM (Figure S10C,F), was no longer detectable at 500 mM (Figure S10G). Confocal imaging of Δ*dltA* co-inoculated with wild type showed sparse GFP-labelled Δ*dltA* cells, relative to mScarlet-I3-labelled wild type, at every root zone examined, including the meristematic zone and lateral root emergence sites where bacteria accumulate most densely (Figure 6C). This uniform contrasts the niche-specificity of Δ*hemAT* and is consistent with a defect in the physical interaction with the root surface.

## Discussion

Understanding the genetic basis of beneficial plant root colonization is not only of fundamental interest but directly relevant to developing more reliable microbial inoculants for sustainable agriculture (14). Yet the mechanisms governing how PGPRs establish themselves on roots remain poorly characterised, particularly in Gram-positive species. Genome-wide Tn-seq screens in Gram-negative *Pseudomonas* have identified motility, biofilm formation, metabolism and immune evasion as key colonization determinants (49, 50). In the Gram-positive PGPR *Bacillus amyloliquefaciens* FZB42, targeted mutagenesis revealed that the regulator DegU governs swarming and biofilm formation, and that *nfrA* and related loci are specifically required for root colonization (51). More recently, inducible CRISPRi systems built for rhizobia (*R. etli*, *S. meliloti*) have enabled targeted essentiality studies in symbiotic bacteria, but CRISPRi has not yet been deployed for pooled, genome-wide forward genetic screens during plant colonization (52, 53). Our CRISPRi-seq screen in *B. subtilis* 3610 addresses the limitations of constitutive loss-of-function approaches, that cannot repress essential genes, as well as of targeted mutagenesis, which does not provide genome scale functional insights (17). Our work thus brings tunable genome-wide functional genomics to a Gram-positive PGPR during active plant colonization and uncovers roles for cell wall remodeling, chemoreceptor signaling and translational reprogramming that prior approaches could not have detected.

One inherent limitation of our screen is that it identifies genes whose loss is more costly on roots than in liquid, without distinguishing whether this reflects a genuine root-specific function or simply a requirement for faster bacterial growth on roots. Faster bacterial growth on roots than in liquid is plausible given their access to root-derived exudates. Indeed, cross-referencing the root-essential hits against the essential genes of the LB control screen reveals that approximately one third encode housekeeping functions. Their appearance thus most likely reflects faster bacterial growth on roots than in liquid, rather than root-specific function. Yet, the remaining two-thirds of genes appear to represent genuinely root-conditional requirements and a growth-corrected comparison (Figure S6, Data S7) maintained significance for all CRISPRi-seq hit taken forward for individual validation. Another limitation is the fact that bacteria were recovered from whole roots, meaning that the fitness costs measured by CRISPRi-seq, as well as the RNA-seq data, do not resolve specific root niches, but represent an average weighted towards the niches carrying most of the population.

An expected finding from the screen was the presence of motility genes in the essential fraction. Here, flagellar proteins and swarming were two of the most significantly enriched functional categories in the CRISPRi-seq GSEA, consistent with the established requirement for *B. subtilis* motility to reach and navigate the root surface (31, 54–56). Interestingly, the appearance of swarming-related loci as essential even in a hydroponic setting suggests that surface motility, not just chemotaxis toward the root, contributes materially to colonization. By contrast, biofilm matrix genes largely failed to score as essential, despite their well-documented role in sustained root association (12). This is expected in a pooled CRISPRi-seq design, where any gene product acting as a public common good will not be detected as a fitness defect because individual bacteria that lack it can exploit these products, such as matrix components secreted by neighbours. A related blind spot concerns inter-bacterial competition. *yukE*, encoding a WXG100-family type VII secretion substrate implicated in *Bacillus* inter-strain antagonism, was among the most strongly induced genes on roots (log_2_FC +2.55) yet expectedly showed no fitness defect in our mono-association CRISPRi-seq screen.

Both “biosynthesis of teichoic acids” and “biosynthesis of isoprenoids” came up as significant SubtiWiki categories in the CRISPRi-seq. Of the 13 genes (11 transcriptional units) in the teichoic-acid category, 8 (4 transcriptional units) were significantly required for root colonization, giving the WTA requirement robust, multi-sgRNA support. The parallel requirement of the isoprenoid/MEP pathway (*dxs*, *ispD*, *ispH*) is also mechanistically consistent, as WTA biosynthesis requires undecaprenyl phosphate as a lipid carrier, assembled from eleven isoprene subunits generated by the MEP pathway. WTAs have additionally been implicated in resistance to cationic antimicrobial peptides (CAMPs) and ROS, suggesting multiple roles for WTAs during root colonization: (i) charge-based attachment scaffold, (ii) shielding from plant-derived antimicrobials, and (iii) protection against ROS/ROS-dependent compounds, generated during plant defense (57, 58). Prior work in *B. velezensis* suggested that the colonization defects of the WTA biosynthetic genes *ggaA* and *gtaB* were due to their requirement for biofilm formation during cucumber root colonization (59). Our screen, where biofilm matrix (possibly due to it being a “public good”) was not detected as essential, prompted us to reveal a more direct, biofilm-independent involvement of teichoic acids in early root attachment. Cell wall remodeling may then be additionally important for adapting to fluctuating osmotic conditions in the root zone (60, 61) and for buffering the bacterial cell against root-derived antimicrobial compounds, ROS, or hydrolytic enzymes (62).

The teichoic acid pathway essentiality in CRISPRi-seq, together with the “cell envelope stress proteins” as an enriched root-upregulated category in RNA-seq points to the root surface as a chemically challenging environment for the bacterial cell envelope. Indeed, *B. subtilis* colonization is known to trigger plant immune priming and systemic resistance in *Arabidopsis* (63), which might initially expose colonizing bacteria to plant-derived antimicrobial peptides, reactive oxygen species, and hydrolytic enzymes at the root surface. Consistently, *B. subtilis* upregulates dedicated stress-response programs upon root association (64). Indeed, we found *B. subtilis* to preferentially colonize root caps and lateral root emergence sites, both shown to display a high intensity of plant immune outputs (65).

WTAs carry a net negative charge that can be modulated by D-alanylation, as well as teichuronic acid under phosphate starvation (66, 67). The strong induction of the Dlt D-alanylation operon suggests that tuning the net surface charge of the cell envelope is a specific response to the root environment. Our direct attachment assays further indicate that part of the requirement for D-alanylation is to render the electrostatics of the bacterial cell wall more compatible with that of plant cell walls. Yet, *dlt* mutants are also hypersensitive to cationic antimicrobial peptides, and transposon libraries grown as mixed populations are depleted of teichoic acid modification mutants, due to their sensitivity to autologous antimicrobials produced by neighbouring cells (68). Based on this, Δ*dltA* might also be more susceptible to root-derived antimicrobials, further adding to their competitive disadvantage on roots.

Among the 14 sgRNAs that significantly improved root fitness when knocked down, both the heme-containing aerotaxis receptor HemAT, and the heme binding protein YwfI were found to be costly. The rhizosphere can be expected to be heterogeneous in oxygen, with hypoxic microniches, particularly in the hydroponic conditions of our assays (69, 70); under these conditions HemAT-guided aerotaxis of wild-type *B. subtilis* could therefore interfere with rapid colonization of nutrient-rich, yet slightly hypoxic root niches, providing HemAT mutants with a competitive advantage. Quantitative image analysis of competitive colonization add a spatial dimension on this effect: the Δ*hemAT*:wild-type fluorescence ratio was approximately tenfold higher at the root cap than at lateral root emergence site, indicating that mutant competitiveness is not uniform along the root, something that is invisible to bulk colonization assays, which integrate over every root niche. This relative advantage of Δ*hemAT* might turn into a disadvantage if exclusively mutant bacteria are present, causing too many bacteria to compete for nutrients in niches with sub-optimal oxygen supply. The root cap is among the most respiratory-active regions of the root and is enveloped in mucilage and sloughing border cells which could plausibly lead to a lower oxygen environment than lateral root emergence site, which are transient, wound-like breaches of the overlying cortex. Direct measurement with oxygen-sensitive probes, or a bacterial hypoxia reporter imaged in the same niches, would be required to substantiate this hypothesis. As mentioned above, wild-type populations on roots contain a higher proportion of long, chained and immotile cells than Δ*hemAT*, leading to an underestimate mutant cell numbers by image-based quantification. Resolving this would require single-cell segmentation or niche-resolved CFU recovery. Nevertheless, the difference in Δ*hemAT* cell size and motility is intriguing, as it suggests that this intracellular aerotaxis receptor has functions beyond strict chemotactic guidance.

An intriguing finding is the concordant induction (RNA-seq) and essentiality (CRISPRi-seq) of *yjbI*, a group-II truncated haemoglobin for nitric oxide (NO) detoxification together with the upregulation of the NO-responsive master regulator *nsrR*. This suggests NO detoxification as a specific requirement for root colonization. The strong induction in an axenic system points to plant-derived NO as a selection pressure for bacterial colonization of root surfaces. While NO is a major component in the bactericidal activity of animal immune cells (71), no similar, oxidative pathway for NO production has been found in plants (72). Yet, a recent publication provides evidence for a new pathway of NO production (73), requiring cell wall-localized peroxidases, superoxide, as well as oxime donors. All these factors are known to be present and active precisely at the sites of *B. subtilis* colonization.

RNA-seq signatures also paint a clear, yet unexpected picture of the carbon utilization of root-colonizing *B. subtilis* on the roots, with upregulation of fructose uptake and catabolic genes, combined with downregulation of the endolevanase *levB* (47) as well as the levansucrase *sacB* (48). This would indicate that root-colonizing *B. subtilis* under our conditions are not obtaining fructose from sucrose or fructose polymers (e.g. levan), but directly take up fructose monomers. The presence of sufficient levels of a preferred carbon source such as fructose would also account for the observed downregulation of genes involved in the utilization of complex plant carbohydrates, such as the pectin degrading *Yes* cluster. Fructose was shown to be a chemo-attractant for *B. subtilis* (74) and was found in root exudates of many species alongside other hexoses, with increased levels in exudates of younger, seedling roots in maize, for example (75, 76). Interestingly, jasmonate mutants of Arabidopsis were found to display increased fructose levels in root exudates, which was correlated with increased representation of *Bacillus* species in complex root microbiota (77). Our data therefore suggests an unexpected carbon source being relevant *for B. subtilis* root colonization. It is currently unclear by which means Arabidopsis seedling roots would exude fructose into the rhizosphere. It has been shown recently that mutants in sugar transporters influence the spatial composition of the root microbiome along the root axis (78) and fructose exporters involved in storage and mobilization of fructose from vacuoles are highly expressed in roots, specifically in root cap cells, coinciding with the preferred site of colonization of *B. subtilis* in our assays (79–81). Moreover, our transcriptional data derive from bacteria recovered from whole roots and therefore average over niches that are not equivalent, as the niche-dependent Δ*hemAT* ratio indicates, and are necessarily weighted towards the sites where most bacteria reside; Fluorescent reporters could resolve such niche differences and reveal whether fructose utilization is a general or a local feature of root colonization. The mechanism of fructose release from root cap cells remains unclear. Plausible routes could be a release through apoptosis of sloughing cap cells, a passive efflux across the plasma membrane or active export by an as-yet-unidentified transporter. This fructose response coupled with TCA cycle induction and ATP synthase upregulation, indicates a state of rapid aerobic growth fueled by root-derived monomeric sugars, a physiological profile distinct from both rich laboratory medium and carbon-limited stationary phase.

This study presents the first genome-wide CRISPRi-seq screen of a Gram-positive PGPR during active plant colonization. It provides a comprehensive view of the requirements for root colonizing *B. subtilis* at single-gene resolution. The screen confirmed expected requirements for motility, while uncovering several unexpected findings: First, a biofilm-independent role of the wall teichoic acid and isoprenoid pathways biosynthesis, with a more specific function for D-alanylation in early attachment and root association; second, a role for monomeric fructose as a dominant carbon source; third, aerotaxis as a competitive disadvantage during colonization and, fourth, the importance of NO detoxification as a discrete colonization requirement, suggestive of an unexpected and little understood role for plant-produced apoplastic NO in root-microbe interactions. More broadly, the CRISPRi-seq library and platform established here is directly transferable to other host plants, as well as soil-based colonization systems, making it a scalable and transferable tool for the rational improvement of bacteria for sustainable agriculture.

## Materials and Methods

### Bacterial and plant materials, growth conditions and bacterial collection

*Bacillus subtilis* NCIB 3610, containing ComIQ12L mutation to enable competence (Bacillus Genetic Stock Center, BGSCID 3A1), and its derivatives were used throughout; all strains are listed in Table S1. Bacteria were revived from frozen stocks on LB agar and grown at 37°C, with liquid cultures shaken at 180 rpm. Chloramphenicol and kanamycin were used at 10 µg/mL and erythromycin at 5 µg/mL. Transformation was performed in MC medium (per litre: 10.70 g K2HPO₄, 5.3 g KH2PO₄, 20 g D-glucose, 1 g casamino acids, 2 g L-glutamate monopotassium salt, 10 mL of 300 mM sodium citrate, 1 mL of 22 mg/mL ferric ammonium citrate), supplemented with tryptophan and/or threonine at 0.5 mg/mL where required: a single colony was grown in 10 mL MC at 37 °C to early stationary phase (OD_600_ 1.0-1.5), and 1 mL of culture mixed with the DNA construct, incubated 40 min at 37 °C and plated on selective LB agar.

*Arabidopsis thaliana* Col-0 (in-house stock) was used throughout; a Col-0 line expressing Lti6b-mTagBFP2 was used for confocal imaging. Seeds were surface-sterilized in 70% (v/v) ethanol containing 0.05% (v/v) Tween 20 for 10 min, rinsed twice in absolute ethanol and air-dried. Seeds were sown at nine per sterile nylon mesh square (2 × 2 cm; Lanz-Anliker AG) on half-strength Murashige and Skoog agar (½ MS; Duchefa) with 0.5 g/L MES, pH 5.7, 0.8% (w/v) agar. Plates were stratified at 4 °C in darkness for 48 h then grown at 23 °C and 65% relative humidity under continuous light for 72 h. Meshes carrying three-day-old seedlings were transferred to 12-well plates containing 6.5 mL per well of liquid ½ MS supplemented with 1 mM sucrose and 0.05 mg/mL L-threonine, at nine seedlings per well, and grown for a further three days, so that plants were six days old at inoculation.

Bacterial cultures were grown in LB from glycerol stocks to mid-exponential phase, harvested by centrifugation and washed in 1× PBS. Suspensions were adjusted to a final OD_600_ of 0.02 in the well and incubated statically under the plant growth conditions above for the times indicated. At harvest, roots were removed and washed by dipping three times in sterile ultrapure water, then transferred to 500 µL of extraction buffer (10 mM MgCl2, 0.01% (v/v) Silwet L-77) and homogenized in a TissueLyser II (Retsch/Qiagen) at 30 Hz for 30 s. The resulting suspension was used for CFU determination or nucleic acid extraction. For the CRISPRi-seq and RNA-seq experiments, the free-swimming population was sampled from 2 mL of the remaining medium after root removal, pelleted at 10,000 × g for 5 min and stored at -80 °C.

### Strain and plasmid construction

Primers and plasmids are listed in Tables S2 and S3. All strains were constructed by homologous recombination in the recipient strain, using natural genetic competence induced at stationary phase upon media depletion. A construct carrying the *Streptococcus pyogenes* dcas9 gene under the Pspank promoter was integrated at the *thrC* locus of *B. subtilis* 3610. A fragment containing the upstream *thrC* sequence and the MLS resistance cassette was amplified with OVL6582/OVL6529, and a second fragment containing *lacI* and the downstream *thrC* sequence with OVL6528/OVL6581, both from pMB002 (modified from pMB001 (82)), a modified pDG1664 derivative carrying Pspank and lacI. The dcas9 fragment was amplified with OVL6530/OVL6531 from pJWV102-PL dCas9 (Addgene #85588). The three fragments were fused by Golden Gate assembly with *Bsa*I. Plasmid pDS22 carries an *Esp*3I-flanked mCherry gene, the dCas9 handle and Illumina adapter sequences under a constitutive promoter (P3), the sgRNA cloning cassette, within *amyE* integration arms, together with a spectinomycin resistance cassette. It was constructed from the backbone of pSG050 (kindly provided by the laboratory of Stephan Gruber, UNIL), a modified pDG3661 vector (GenBank AY618310) lacking *lacZ*. pSG050 was linearized with OVL6453/OVL6454 and the sgRNA cloning cassette amplified with OVL6455/OVL6456; the two fragments were joined by Golden Gate assembly with *Aar*I.

Deletion mutants were constructed from the *B. subtilis* 168 BKK kanamycin knockout collection (83). For each target, the deletion allele was amplified with approximately 1 kb of flanking homology on each side, introduced into *B. subtilis* 3610 by natural transformation and homologous recombination, and selected on kanamycin. Deletions were verified by colony PCR across both junctions and by sequencing.

Deletion mutants were not complemented in trans. Instead, each phenotype was attributed using the CRISPRi-seq data (Data S4), as follows. A deletion phenotype may arise from a secondary or suppressor mutation, from the cassette itself, or from polarity on downstream genes. The first two are addressed by concordance with CRISPRi-seq: deletion and dCas9 knockdown are distinct interferences in independently constructed strains, and are not expected to share a secondary mutation, a cassette artefact or an off-target. Polarity, however, affects both perturbations, so it was assessed directly from the sgRNAs targeting the genes lying downstream of each deletion within the same transcriptional unit, these being the only genes a polar effect can reach. Throughout, significance refers to the root-colonization hit criteria (|log_2_FC| ≥ 1, padj < 0.05). *hemAT*, *lexA*, *hag* and *rghRA* are monocistronic. Downstream of *reoM/yrzL* in *reoM-yrrK-yrzB*, neither *yrrK* nor *yrzB* met the hit criteria, and the effect of *yrrK* was in the opposite, mildly costly direction (+0.63). In *yopM-yopN-yopO-yopP-yopQ-mrpR*, the only gene downstream of *yopQ* is *mrpR/yopR*, which was unaffected (−0.18, ns). In *ecsA-ecsB-ecsC*, the downstream *ecsC* was far less depleted than *ecsB* itself (−0.40, ns, against -2.60). In none of these cases can polarity account for the mutant phenotype. Three loci are not resolved to the gene. In *cdaA-cdaR-glmM*, *cdaR* was depleted as strongly as *cdaA* (−2.76 against -2.78) and *glmM* was also required (−1.77), so the requirement is attributed to the operon and the contributions of c-di-AMP and peptidoglycan precursor synthesis cannot be separated. In the *fla/che* operon, all three genes downstream of *cheC* were significantly depleted (*cheD* -0.79, *sigD* -1.34, *ylxL*-0.63; *cheD* and *ylxL* below the effect-size threshold), with the sigma factor *sigD* showing the strongest effect, so the Δ*cheC* defect is attributed to the locus rather than to CheC specifically. In *tagD-tagE-tagF-tagG-tagH*, the genes downstream of *tagE* were all significantly required (*tagF* -1.58, *tagG* -2.42, *tagH* -1.30), consistent with the multi-sgRNA support for wall teichoic acid biosynthesis as a whole. The Δ*tagE* phenotype is therefore interpreted as evidence for a requirement for WTA biosynthesis rather than for TagE specifically.

### Construction of the pooled CRISPRi-seq library

The *B. subtilis* sgRNA library (data S14) was designed as previously described (16) using the pipeline at https://github.com/veeninglab/CRISPRi-seq and the genome of *B. subtilis* 168 strain VL4311 (GenBank CP171153). Each sgRNA carries Esp3I flanking regions for Golden Gate cloning, strain-specific priming sequences for amplification of the *B. subtilis* sub-library, and universal sequences for amplification of the whole pangenome library. The sub-library was amplified from the synthetic pangenome pool (Veening lab resource), and cloned into linearized pDS22 by Golden Gate assembly with Esp3I (84). Correct assembly replaces *mCherry*, so clones carrying an sgRNA are white rather than red. Approximately 250,000 *E. coli* colonies were obtained and pooled, none red, giving >50-fold theoretical coverage of the 4,326 sgRNAs in the library. The pooled plasmid library was transformed into *B. subtilis* 3610 carrying *thrC*::P*spank*-*dCas9*-*ery*, integrating the sgRNA cassette at *amyE*; approximately 250,000 colonies were scraped and stored at -80 °C as the CRISPRi library. For all experiments the library was inoculated into LB to a starting OD_600_ of 0.3 and grown overnight at 37 °C with shaking. Library representation was verified by amplicon sequencing. Individual sgRNA strains targeting two essential genes (*murF*, *dnaA*) and one non-essential gene (*ganA*) were constructed as above, cloning a single sgRNA spacer into pDS22. Strains were inoculated to OD_600_ 0.05 in plant exudate medium supplemented with 15 mM sucrose and grown at 37 °C, 180 rpm, with or without 1 mM IPTG; OD_600_ was recorded at 8 h and 24 h. Three biological replicates were performed per strain and condition. Plant exudate medium was conditioned medium collected from Col-0 seedlings grown for one week in the hydroponic system above.

### CRISPRi-seq screens

For the LB screen, the CRISPRi library was grown overnight in LB at 37 °C with shaking, diluted to OD_600_ 0.02 in fresh LB with or without 1 mM IPTG and grown for approximately six generations, then re-diluted to OD_600_ 0.02 under the same conditions for a further six generations, giving approximately twelve generations of induction. Cells were harvested (4,000 × g, 15 min), washed twice in PBS and stored at -80 °C. Four biological replicates were performed per condition. For the plant root screen, the library was pre-induced in LB with or without 1 mM IPTG for approximately four generations, harvested, washed in 1× PBS and inoculated into the hydroponic system at a final OD_600_ of 0.02. Induced wells received a further 2 mM IPTG at inoculation and again at 6 h to maintain knockdown. At 24 h, root-attached and free-swimming populations were harvested separately as described above and stored at -80 °C. Four biological replicates were performed per condition, each comprising 108 seedlings (twelve wells of nine seedlings) harvested and pooled.

### DNA and RNA extractions

Genomic DNA was extracted from cell pellets using the Wizard Genomic DNA Purification Kit (Promega) according to the manufacturer’s instructions, with a lysozyme pre-treatment (10 µL of 100 µg/µL lysozyme in 450 µL of 50 mM EDTA, 20 min at 37 °C) preceding lysis and an RNase A step (4 µg/µL, 15 min at 37 °C). DNA was rehydrated in 200 µL of TE buffer at 65 °C and stored at -20 °C. Total RNA was isolated from *B. subtilis* cell pellets, that were treated RNAprotect (Qiagen), using the QIAwave RNA Mini Kit (Qiagen) with a modified lysis step (85): pellets were resuspended in 200 µL of lysozyme solution (20 mM Tris-HCl pH 8, 2 mM EDTA, 1.2% (v/v) Triton X-100, 20 mg/mL lysozyme) and incubated at 37 °C for 30 min before processing according to the manufacturer’s instructions, including the on-column DNase treatment. RNA was eluted in 30 µL of RNase-free water. Four biological replicates were prepared per condition, each comprising 108 seedlings harvested and pooled. Nucleic acid quality and quantity were assessed with a NanoDrop (Thermo Fisher) and a Fragment Analyzer (Agilent Technologies).’

### Sequencing library preparation and sequencing

For CRISPRi-seq, the sgRNA amplicon was generated by PCR with oligonucleotides complementary to the Read 1 and Read 2 Illumina adapter sequences carried within the chromosomally integrated sgRNA cassette; P5 and P7 adapters were added in the same reaction and libraries dual indexed with IDT10 UDI index pairs. For RNA-seq, ribosomal RNA was depleted with RiboCop rRNA Depletion Kits (Lexogen) for bacteria and for plant, applied as a combined probe pool to remove both bacterial and residual *Arabidopsis* rRNA. Depleted RNA was converted to libraries with the Watchmaker RNA Library Prep Kit (Watchmaker Genomics) and dual indexed with IDT10 UDI index pairs. Eight libraries, four root and four liquid, were pooled and sequenced on an AVITI system (Element Biosciences) in 150-base single-end mode at the Lausanne Genomics Technologies Facility, each library across two lanes. Base calling and demultiplexing used Bases2Fastq v2.10.

### CRISPRi-seq analysis

Raw FASTQ files were processed with 2FAST2Q v2.8.1 to obtain per-sgRNA read counts, using a start position of 54, a feature length of 20 bp, a maximum of two mismatches against the library spacer sequences and a minimum Phred quality threshold of 10. Count matrices were analysed in R v4.5.3 with DESeq2 v1.50.2. In each analysis, sgRNAs with fewer than 10 reads summed across all samples were removed, size factors and dispersions were estimated from the data, and significance was assessed by Wald test at an FDR of α = 0.05. Log2 fold-changes were shrunk with apeglm v1.32.0 except where stated.

Counts from the LB screen were modelled with the design ∼ IPTG (reference 0 µM), testing the coefficient IPTG_1mM_vs_0uM against a null of no effect (lfcThreshold = 0); sgRNAs with adjusted *p* < 0.05 and absolute shrunken log2 fold-change ≥ 1 were considered depleted. Counts from the pre-induction experiment used the same design but were tested against a non-zero null (lfcThreshold = 1), testing the hypothesis that |log_2_FC| ≤ 1, with unshrunken log2 fold-changes. No sgRNA was significantly depleted, indicating uniform library representation at inoculation. Counts from the colonization experiment were modelled with the design ∼ background * IPTG, where background has two levels (3610_liquid, the library recovered from the medium surrounding the seedlings, and 3610_root, the library recovered from the roots of the same wells), with 3610_liquid and 0 µM IPTG as references. The interaction term background3610_root.IPTG1mM, the difference between the IPTG effects in each background, was used to identify sgRNAs whose knockdown imposes a greater fitness cost during root colonization than in liquid, and was tested against a null of no interaction (lfcThreshold = 0). sgRNAs with adjusted *p* < 0.05 and absolute shrunken interaction log2 fold-change ≥ 1 were considered hits.

The interaction contrast compares induction effects per experiment, assuming equal numbers of generations in the two compartments. Since viable counts were not obtained, the ratio was estimated from the library: the 599 sgRNAs significant (FDR < 0.05) and costly (|log_2_FC| ≤ -1) in the LB screen (Data S1) were taken as calibrators whose cost should be dominated by growth rate, and their shrunken effects in liquid and on roots related by Deming (errors-in-variables) regression through the origin, with the error-variance ratio set to the ratio of mean squared standard errors and a 95% CI from 500 bootstrap resamples of sgRNAs (β = 1.44, 95% CI 1.38-1.51; 1.49 fitting all sgRNAs, 1.38 with a free intercept). Cost per generation was then tested as the contrast (root effect) - β × (liquid effect) in the same DESeq2 model, leaving design, dispersions, Wald test and Benjamini-Hochberg adjustment unchanged. Because apeglm shrinks single coefficients and not arbitrary contrasts, this contrast and the β = 1 contrast were both shrunk with ashr. sgRNAs with FDR < 0.05 and |log_2_FC| ≥ 1 were called; counts were stable across the confidence interval of β (126 at 1.44, 130 at 1.38, 122 at 1.51). Gene-level aggregation and GSEA followed the procedures described above.

### RNAseq analysis

Libraries were prepared from four biological replicates per condition. Two were excluded before any differential expression testing. Root_C was a technical replicate of Root_A rather than an independent biological sample (Spearman r = 0.999 on rlog-transformed counts, against 0.985 for the next-closest pair of libraries). Liquid_B was excluded on global-similarity grounds: it was the only library whose mean correlation to its own condition group was lower than to the opposite group (−0.011, against +0.009 to +0.043 for every other library), and it fell within the root range on the first principal component (PC1 = -8.9, root range -11.6 to -7.3, retained liquid libraries +3.2 to +24.3). Its single closest library nonetheless remained Liquid_A, so the anomaly is one of position between the groups rather than membership of the root group. Differential expression was therefore performed on three biological replicates per condition.

To characterise the Liquid_B anomaly, we subsequently scored all eight libraries on a root-versus-liquid signature comprising the 50 most strongly root-enriched and 50 most strongly liquid-enriched genes from the final three-versus-three contrast (padj < 0.01). Because that contrast excludes Liquid_B, the library is an independent test point. Projected onto a liquid(0)– root(1) axis anchored on the six retained libraries, the four root libraries scored 0.92 to 1.09 and the three retained liquid libraries -0.24, 0.01 and 0.24, whereas Liquid_B scored 0.60, with 81% of the 37 markers whose root–liquid separation exceeded 1.5 rlog units falling on the root side of the axis midpoint and a unimodal per-marker distribution (interquartile range 0.59 to 0.85). The shift was graded across essentially all markers rather than confined to a subset of genes, and was not attributable to library composition: marker score scales with the fraction of reads assigned to bacterial coding sequence within both conditions (+0.024 per percentage point, fitted on the six retained libraries), and after adjusting for this Liquid_B remained 3.8 residual standard deviations above the expectation for a liquid library and 7.4 below the expectation for a root library. Simple mislabelling is also inconsistent with the data, since 94.0% of Liquid_B reads aligned to the *B. subtilis* genome against 22.3 to 55.0% for the root libraries, and its residual rRNA fraction was the lowest of the eight (1.30%). Partial mixing or carry-over during preparation, or detached root-associated cells present in the liquid fraction, are more consistent with the graded pattern. Because no condition label could be assigned to this library, it was excluded from the two-group model. To confirm that this exclusion does not drive our conclusions, differential expression was repeated including all eight libraries. Log2 fold changes were highly concordant with the three-versus-three analysis (Pearson r = 0.942 across 4,290 genes), with 875 genes significant in both. Per-library metrics, clustering and marker diagnostics are given in Figure S11 and Data S15.

Reads from the two sequencing lanes were concatenated per library, then adapter-trimmed and quality-filtered with fastp v0.23.2 using default thresholds. Filtered reads were aligned to the *B. subtilis* NCIB 3610 chromosome (CP020102.1) together with the pBS32 plasmid (CP020103.1) using BWA-MEM2 v2.2.1 with default parameters, sorted and indexed with SAMtools v1.21, and assigned to genes with featureCounts v2.0.3 restricting features to coding sequences (−t CDS) and grouping by locus tag (−g locus_tag). pBS32 is absent from *B. subtilis* 168 and therefore unrepresented in the CRISPRi library, so plasmid-encoded genes are retained in the RNA-seq differential expression results but absent from the integrated analyses. The count matrix was analysed with DESeq2 v1.50.2. Genes with fewer than 10 reads summed across all samples were removed. Counts were modelled with the design ∼ condition (reference Liquid) and Wald tests used at an FDR of α = 0.05. Log2 fold-changes are reported as unshrunken maximum-likelihood estimates; genes with adjusted *p* < 0.05 and absolute log2 fold-change ≥ 1 were considered differentially expressed. Regularized-log transformed counts (blind = FALSE) were used for principal component analysis, sample-to-sample correlation and heatmaps. Gene-level annotation used rtracklayer v1.70.1 from the genome GFF3 file.

### Gene set enrichment analysis

Gene sets were constructed from the functional category annotations of SubtiWiki (33), restricted to 5–500 members. Category labels differing only by a trailing gene count were merged and duplicate gene sets collapsed to a single representative. Gene identifiers were matched to SubtiWiki entries through both canonical names and the synonym field, so that loci carrying legacy names in the sgRNA library annotation were retained. The same gene-set construction was applied to both datasets. Enrichment was tested with fgsea v1.36.2 using fgseaMultilevel with eps = 0 and nPermSimple = 100000. For CRISPRi-seq, genes were ranked by the negative of their median shrunken interaction log2 fold-change, so that a positive normalized enrichment score identifies categories enriched among genes required for root colonization; SubtiWiki regulon annotations were additionally included as gene sets. For RNA-seq, genes were ranked by sign(log2 fold-change) × -log₁₀(adjusted *p*), so that a positive score identifies categories enriched among genes induced on roots. Categories with adjusted *p* < 0.05 were considered significant. Random seeds are recorded in the deposited code.

### Bacterial quantification after root colonization

Nine roots per well were homogenized in a common volume of 500 µL as described above. A 40 µL aliquot was taken into a 1-in-5 serial dilution series in sterile water, and 20 µL of each dilution spotted onto LB agar containing the appropriate antibiotic. Plates were incubated overnight at 37°C and colonies counted at a dilution giving a countable number. Colony-forming units per millilitre of homogenate were calculated as CFU/mL = colonies × 5ⁿ / 0.02, where n is the dilution step, the first dilution corresponding to a 5-fold dilution of the homogenate. Colonization was expressed per root as CFU/root = CFU/mL × 0.5 mL / 9.

For individual colonization assays, each strain was inoculated alone into the hydroponic system at a final OD_600_ of 0.02 and roots harvested at 24 h. For competition assays, wild-type *B. subtilis* marked with a chloramphenicol cassette and the mutant marked with a kanamycin cassette were grown separately, mixed 1:1 and co-inoculated at a final OD_600_ of 0.01 per strain. Roots were harvested at 24 h and both populations enumerated in parallel from the same homogenate on LB agar containing either chloramphenicol or kanamycin.

For early attachment assays, Col-0 seedlings were grown vertically on ½ MS agar for six days and roots inoculated with 10 µL of bacterial suspension at OD_600_ 0.02. After 30 min or 2 h, roots were harvested, washed with sterile water and processed as above. Attachment was expressed as a percentage of the mean wild-type value within each timepoint; because normalization is performed within a timepoint, the dilution factor and per-root normalization cancel and do not affect the reported percentages. Six biological replicates were performed per group, each comprising nine pooled seedlings.

### Inert attachment assay

Assays were performed in base assay buffer (BAB: 10 mM sodium phosphate pH 7.0, 50 mM NaCl). For the ionic strength series, NaCl was added to 10 mM sodium phosphate pH 7.0 to give 0, 50 or 500 mM, corresponding to ionic strengths of 18, 68 and 518 mM and Debye lengths of 2.29, 1.17 and 0.42 nm; the phosphate base contributes 17.7 mM, so the 0 mM condition denotes no added NaCl rather than zero ionic strength. Each condition used its own buffer for cell resuspension, washing and recovery. Tissue-culture-treated flat-bottom 96-well plates were coated with poly-L-lysine (PLL, MW 70,000–150,000 Da; 0.1 mg/mL in ultrapure water, 100 µL per well, 1 h at room temperature) or polygalacturonic acid (PGA) sodium salt (2 mg/mL in 10 mM MES pH 5.5, 100 µL per well, overnight at 4 °C), then washed with ultrapure water and air-dried; uncoated wells served as a moderately negative reference surface. PGA coating was confirmed in separate wells by Alcian Blue staining. All wells were then blocked with 200 µL of 0.5% (w/v) BSA in BAB for 30 min at room temperature, which was aspirated without further washing.

Bacteria were grown to mid-exponential phase, washed three times in the relevant assay buffer and diluted to OD_600_ 0.05; input suspensions were plated to confirm equivalent cell numbers between strains. Wells received 100 µL of suspension, or buffer alone as a no-bacteria control, and were incubated statically at 25°C for 30 min or 2 h. Non-adherent cells were removed by three wash cycles, each comprising aspiration of 90 µL, gentle addition of 150 µL buffer down the side of the well, a 30 s pause and aspiration. Adherent bacteria were recovered into 100 µL of buffer by pipetting ten times. For the salt shock and return condition, cells were pre-incubated for 30 min in 10 mM sodium phosphate pH 7.0 containing 500 mM NaCl, returned to BAB and assayed on PGA in BAB. Recovered suspensions were diluted and plated as above, using the corresponding assay buffer as diluent. Attachment was expressed as a percentage of the mean wild-type value on the same surface, in the same buffer and at the same timepoint; absolute CFU/mL were used for comparisons between buffers. Six independent biological replicates were performed per group.

### Confocal microscopy

Imaging was performed on a Leica Stellaris 5 confocal microscope using a 63X water immersion objective. mScarlet-I3-labeled wild-type *B. subtilis*, inoculated either alone or together with one of four GFP-labeled mutants was imaged on Arabidopsis roots expressing mTagBFP2 in the plasma membrane. For the early differentiation zone, differentiation zone, and the lateral root emergence zone, a single field of 512 × 512 pixels was acquired. For the root tip region, a mosaic of 12 tiles (512 × 512 pixels each) was acquired. Images were acquired using two sequential detection tracks. Track 1 captured mTagBFP2 and mScarlet-I3 (Ex. 405 nm and 561 nm; Em. 430-480 nm and 567-619 nm, respectively). Track 2 captured GFP (Ex. 488 nm; Em. 500-540 nm). A z-stack of 29 slices (2 µm step size) was acquired, spanning from the root surface to a depth of 55 µm. Images of wild-type *B. subtilis* co-inoculated with Δ*tagE,* Δ*teoM,* Δ*dltA and* Δ*hemAT B. subtilis* were spectrally unmixed in Fiji/ImageJ2 (version 2.16.0/1.54p) to correct a spectral overlap between mScarlet-I3 and GFP. A maximum-intensity z-projection was generated on composite images. Reference emission fingerprints for mTagBFP2, mScarlet-I3 and GFP were built once, from single-fluorophore control samples, using the spectral-unmixing tools of the Jay Unruh plugin suite (Stowers Institute) and saved as a combined, normalized reference file. Each maximum-intensity composite was then unmixed against this reference using the linear unmixing jru v2 plugin (Fiji, Jay Unruh/Stowers Institute plugin suite). The three unmixed channels were false colored (GFP, green; mScarlet-I3, magenta; mTagBFP2, cyan) and merged into a single composite overlay.

Images of the GFP-expressing HemAT mutant and mScarletI3-expressing wild type were analyzed in Fiji/ImageJ. Images were split into single channels, and maximum intensity projections were then generated for each channel. The root cap (RC) and lateral root (LR) niches were analyzed separately. To exclude WT bacteria from the GFP signal, the mScarlet channel was thresholded at a fixed intensity of 64, the resulting selection was restored onto the corresponding GFP image, and the pixels within it were set to zero. This step removes GFP pixels overlapping WT signal. GFP and mScarlet pixels were then counted after thresholding each channel. Because background noise differs between niches, GFP was thresholded at 29 in RC and 170 in LR, while mScarlet was thresholded at 64 in both niches. Thresholds were determined by visual inspection and were fixed before the ratios were calculated. For each image, the relative abundance of the HemAT mutant was calculated as the ratio of GFP-positive pixels to mScarlet-positive pixels. RC and LR images were acquired from the same roots, and only roots with both an RC and an LR image were retained for analysis. n refers to the number of roots (n = 10 roots from three independent experiments). Niches were compared by exact two-sided Mann-Whitney U test.

## Supplementary information

Supplementary Information is available for this paper. Supplementary figures and tables are provided in a separate Word document and supplementary data is provided in a separate Excel document.

## Data availability

Processed data are provided as a Supplementary Data Excel file. CRISPRi-seq data generated in this study have been deposited to ArrayExpress (BioStudies) under accession number E-MTAB-16785. RNA-seq data generated in this study have been deposited to ArrayExpress (BioStudies) under accession number E-MTAB-16787. Code used for analysis of the CRISPRi-seq and RNA-seq datasets have been uploaded to Zenodo with the DOI 10.5281/zenodo.21885656.

## Author contributions

Conceptualization, V.M.; Methodology, V.M., J.C.A., D.S., J.-W.V., and N.G.; Software and Formal Analysis, V.M.; Investigation, V.M. and W.A.; Resources, J.C.A., V.D.T., D.S., J.-W.V., and N.G.; Data Curation, V.M.; Visualization, V.M. and H.-H.T.; Writing, V.M., J.C.A., N.G., J.-W.V., W.A. and H.-H.T.; Supervision, N.G. and J.-W.V.; Funding Acquisition, N.G.

## Declaration of interests

The authors declare no competing interests.

## Acknowledgements

The authors would like to thank the Genomics Technology Facility (GTF) at the University of Lausanne (UNIL) for their support with CRISPRi and RNA library preparation and sequencing. Work in the lab of N.G. was supported by the ERC AdG ROOBABAA (101020794) and the SNSF grants 10004646 and 10002702. Work in the lab of J.W.V. was supported by SNSF grants 320030-231669, 320030-236203 and NCCR 51NF40_180541. We would also like to thank Mathilde Ferry for her help in the root glycerol vs sucrose experiments.

The funders had no role in study design, data collection and analysis, decision to publish or preparation of the manuscript. Portions of the introduction, results, and discussion were edited for clarity and language with the assistance of Claude (Anthropic), an AI language model. All scientific content, interpretations, and conclusions are the authors’ own.

## Notes

https://www.ebi.ac.uk/biostudies/arrayexpress/studies/E-MTAB-16785

https://www.ebi.ac.uk/biostudies/arrayexpress/studies/E-MTAB-16787

### Competing Interest Statement

The authors have declared no competing interest.

### Summary of Updates

This version adds an analysis correcting the root versus liquid comparison for differences in generation number between the two populations. A new supplementary figure and a new supplementary data file present this analysis. A new paragraph in the CRISPRi-seq analysis methods describes the calibration and the contrast. The Results and Discussion have been updated to report the corrected analysis and its consequences for interpretation of the cell envelope hits. Additionally, new methods paragraph explains how deletion phenotypes were attributed, the methods heading Genomic extractions has been changed to DNA and RNA extractions, confocal images of tagE KO condition and hemAT KO condition in Figure 3 were swapped in F1 and are now corrected, and the spelling of one author name has been corrected.

## References

1. Trivedi P, Leach JE, Tringe SG, Sa T, Singh BK. 2020. Plant–microbiome interactions: from community assembly to plant health. Nat Rev Microbiol 18:607–621.

2. Blake C, Christensen MN, Kovács ÁT. 2021. Molecular Aspects of Plant Growth Promotion and Protection by Bacillus subtilis. MPMI 34:15–25.

3. Mahapatra S, Yadav R, Ramakrishna W. 2022. Bacillus subtilis impact on plant growth, soil health and environment: Dr. Jekyll and Mr. Hyde. Journal of Applied Microbiology 132:3543– 3562.

4. Su Y, Liu C, Fang H, Zhang D. 2020. Bacillus subtilis: a universal cell factory for industry, agriculture, biomaterials and medicine. Microbial Cell Factories 19:173.

5. He C, Gao T, Wang X, Chen R, Gao H, Liu H. 2025. Co-inoculation of Bacillus subtilis and Bradyrhizobium liaoningense increased soybean yield and improved soil bacterial community composition in coastal saline-alkali land. Front Plant Sci 16:1677763.

6. Singh RK, Singh P, Li H-B, Song Q-Q, Guo D-J, Solanki MK, Verma KK, Malviya MK, Song X-P, Lakshmanan P, Yang L-T, Li Y-R. 2020. Diversity of nitrogen-fixing rhizobacteria associated with sugarcane: a comprehensive study of plant-microbe interactions for growth enhancement in Saccharum spp. BMC Plant Biology 20:220.

7. Jensen CNG, Pang JKY, Gottardi M, Kračun SK, Svendsen BA, Nielsen KF, Kovács ÁT, Moelbak L, Fimognari L, Husted S, Schulz A. 2024. Bacillus subtilis promotes plant phosphorus (P) acquisition through P solubilization and stimulation of root and root hair growth. Physiol Plant 176:e14338.

8. Ryu C-M, Farag MA, Hu C-H, Reddy MS, Wei H-X, Paré PW, Kloepper JW. 2003. Bacterial volatiles promote growth in Arabidopsis. Proceedings of the National Academy of Sciences 100:4927–4932.

9. Kloepper JW, Ryu C-M, Zhang S. 2004. Induced Systemic Resistance and Promotion of Plant Growth by *Bacillus* spp. Phytopathology® 94:1259–1266.

10. Beauregard PB, Chai Y, Vlamakis H, Losick R, Kolter R. 2013. Bacillus subtilis biofilm induction by plant polysaccharides. Proc Natl Acad Sci U S A 110:E1621–E1630.

11. Vlamakis H, Aguilar C, Losick R, Kolter R. 2008. Control of cell fate by the formation of an architecturally complex bacterial community. Genes Dev 22:945–953.

12. Vlamakis H, Chai Y, Beauregard P, Losick R, Kolter R. 2013. Sticking together: building a biofilm the Bacillus subtilis way. Nat Rev Microbiol 11:157–168.

13. Tian T, Sun B, Shi H, Gao T, He Y, Li Y, Liu Y, Li X, Zhang L, Li S, Wang Q, Chai Y. 2021. Sucrose triggers a novel signaling cascade promoting Bacillus subtilis rhizosphere colonization. ISME J 15:2723–2737.

14. de Andrade LA, Santos CHB, Frezarin ET, Sales LR, Rigobelo EC. 2023. Plant Growth-Promoting Rhizobacteria for Sustainable Agricultural Production. Microorganisms 11:1088.

15. Sun L, Zheng P, Sun J, Wendisch VF, Wang Y. 2023. Genome-scale CRISPRi screening: A powerful tool in engineering microbiology. Engineering Microbiology 3:100089.

16. de Bakker V, Liu X, Bravo AM, Veening J-W. 2022. CRISPRi-seq for genome-wide fitness quantification in bacteria. Nat Protoc 17:252–281.

17. Cain AK, Barquist L, Goodman AL, Paulsen IT, Parkhill J, van Opijnen T. 2020. A decade of advances in transposon-insertion sequencing. Nat Rev Genet 21:526–540.

18. Cui L, Vigouroux A, Rousset F, Varet H, Khanna V, Bikard D. 2018. A CRISPRi screen in E. coli reveals sequence-specific toxicity of dCas9. Nat Commun 9:1912.

19. Liu X, de Bakker V, Heggenhougen MV, Mårli MT, Frøynes AH, Salehian Z, Porcellato D, Morales Angeles D, Veening J-W, Kjos M. 2024. Genome-wide CRISPRi screens for high-throughput fitness quantification and identification of determinants for dalbavancin susceptibility in Staphylococcus aureus. mSystems 9:e01289–23.

20. Liu X, Kimmey JM, Matarazzo L, de Bakker V, Van Maele L, Sirard J-C, Nizet V, Veening J- W. 2021. Exploration of Bacterial Bottlenecks and Streptococcus pneumoniae Pathogenesis by CRISPRi-Seq. Cell Host Microbe 29:107–120.e6.

21. Wang X, Jowsey WJ, Cheung C-Y, Smart CJ, Klaus HR, Seeto NE, Waller NJ, Chrisp MT, Peterson AL, Ofori-Anyinam B, Strong E, Nijagal B, West NP, Yang JH, Fineran PC, Cook GM, Jackson SA, McNeil MB. 2024. Whole genome CRISPRi screening identifies druggable vulnerabilities in an isoniazid resistant strain of Mycobacterium tuberculosis. Nat Commun 15:9791.

22. Gil-Campillo C, Mignolet J, Pedro AD-S, Rapún-Araiz B, Janssen AB, Bakker V de, Veening J-W, Garmendia J. 2025. CRISPRi-seq in Haemophilus influenzae reveals genome-wide and medium-specific growth determinants. PLOS Pathogens 21:e1013650.

23. Zhu X, Luo H, Yu X, Lv H, Su L, Zhang K, Wu J. 2024. Genome-Wide CRISPRi Screening of Key Genes for Recombinant Protein Expression in Bacillus Subtilis. Advanced Science 11:2404313.

24. Kim LM, Koo B-M, Burkhart HN, Vollmer D, Vollmer W, Todor H, Gross CA. 2026. The phenotypic landscape of the model firmicute Bacillus subtilis. Proceedings of the National Academy of Sciences 123:e2619440123.

25. 2024. Shedding light on bacteria–host interactions with the aid of TnSeq approaches. mBio 15.

26. McLoon AL, Guttenplan SB, Kearns DB, Kolter R, Losick R. 2011. Tracing the Domestication of a Biofilm-Forming Bacterium. J Bacteriol 193:2027–2034.

27. Burton AT, Kearns DB. 2020. The Large pBS32/pLS32 Plasmid of Ancestral Bacillus subtilis. J Bacteriol 202:e00290–20.

28. Peters JM, Colavin A, Shi H, Czarny TL, Larson MH, Wong S, Hawkins JS, Lu CHS, Koo B- M, Marta E, Shiver AL, Whitehead EH, Weissman JS, Brown ED, Qi LS, Huang KC, Gross CA. 2016. A Comprehensive, CRISPR-based Functional Analysis of Essential Genes in Bacteria. Cell 165:1493–1506.

29. Zhou F, Emonet A, Dénervaud Tendon V, Marhavy P, Wu D, Lahaye T, Geldner N. 2020. Co-incidence of Damage and Microbial Patterns Controls Localized Immune Responses in Roots. Cell 180:440–453.e18.

30. Charron-Lamoureux V, Beauregard PB. 2019. Arabidopsis thaliana Seedlings Influence Bacillus subtilis Spore Formation. MPMI 32:1188–1195.

31. Allard-Massicotte R, Tessier L, Lécuyer F, Lakshmanan V, Lucier J-F, Garneau D, Caudwell L, Vlamakis H, Bais HP, Beauregard PB. 2016. Bacillus subtilis Early Colonization of Arabidopsis thaliana Roots Involves Multiple Chemotaxis Receptors. mBio 7:10.1128/mbio.01664-16.

32. Hu J, Zhang Y, Wang J, Zhou Y. 2014. Glycerol Affects Root Development through Regulation of Multiple Pathways in Arabidopsis. PLOS ONE 9:e86269.

33. Elfmann C, Dumann V, van den Berg T, Stülke J. 2025. A new framework for SubtiWiki, the database for the model organism Bacillus subtilis. Nucleic Acids Res 53:D864–D870.

34. Hu G, Wang Y, Blake C, Nordgaard M, Liu X, Wang B, Kovács ÁT. 2023. Parallel genetic adaptation of Bacillus subtilis to different plant species. Microb Genom 9:mgen001064.

35. Sanchez S, Snider EV, Wang X, Kearns DB. 2022. Identification of Genes Required for Swarming Motility in Bacillus subtilis Using Transposon Mutagenesis and High-Throughput Sequencing (TnSeq). J Bacteriol 204:e0008922.

36. Kirby JR, Kristich CJ, Saulmon MM, Zimmer MA, Garrity LF, Zhulin IB, Ordal GW. 2001. CheC is related to the family of flagellar switch proteins and acts independently from CheD to control chemotaxis in Bacillus subtilis. Mol Microbiol 42:573–585.

37. Hölscher T, Bartels B, Lin Y-C, Gallegos-Monterrosa R, Price-Whelan A, Kolter R, Dietrich LEP, Kovács ÁT. 2015. Motility, chemotaxis and aerotaxis contribute to competitiveness during bacterial pellicle biofilm development. J Mol Biol 427:3695–3708.

38. Raymond-Denise A, Guillen N. 1991. Identification of dinR, a DNA damage-inducible regulator gene of Bacillus subtilis. J Bacteriol 173:7084–7091.

39. Gundlach J, Mehne FMP, Herzberg C, Kampf J, Valerius O, Kaever V, Stülke J. 2015. An Essential Poison: Synthesis and Degradation of Cyclic Di-AMP in Bacillus subtilis. J Bacteriol 197:3265–3274.

40. Allison SE, D’Elia MA, Arar S, Monteiro MA, Brown ED. 2011. Studies of the Genetics, Function, and Kinetic Mechanism of TagE, the Wall Teichoic Acid Glycosyltransferase in Bacillus subtilis 168 *. Journal of Biological Chemistry 286:23708–23716.

41. Yanouri A, Daniel RA, Errington J, Buchanan CE. 1993. Cloning and sequencing of the cell division gene pbpB, which encodes penicillin-binding protein 2B in Bacillus subtilis. J Bacteriol 175:7604–7616.

42. Wamp S, Rutter ZJ, Rismondo J, Jennings CE, Möller L, Lewis RJ, Halbedel S. 2020. PrkA controls peptidoglycan biosynthesis through the essential phosphorylation of ReoM. Elife 9:e56048.

43. Meeske AJ, Rodrigues CDA, Brady J, Lim HC, Bernhardt TG, Rudner DZ. 2016. High-Throughput Genetic Screens Identify a Large and Diverse Collection of New Sporulation Genes in Bacillus subtilis. PLoS Biology 14:e1002341.

44. Hayashi K, Kensuke T, Kobayashi K, Ogasawara N, Ogura M. 2006. Bacillus subtilis RghR (YvaN) represses rapG and rapH, which encode inhibitors of expression of the srfA operon. Mol Microbiol 59:1714–1729.

45. Zhang W, Olson JS, Phillips GN. 2005. Biophysical and kinetic characterization of HemAT, an aerotaxis receptor from Bacillus subtilis. Biophys J 88:2801–2814.

46. Martin-Verstraete I, Débarbouillé M, Klier A, Rapoport G. 1990. Levanase operon of *Bacillus subtilis* includes a fructose-specific phosphotransferase system regulating the expression of the operon. Journal of Molecular Biology 214:657–671.

47. Pereira Y, Petit-Glatron MF, Chambert R. 2001. yveB, Encoding endolevanase LevB, is part of the sacB-yveB-yveA levansucrase tricistronic operon in Bacillus subtilis. Microbiology (Reading) 147:3413–3419.

48. Méndez-Lorenzo L, Porras-Domínguez JR, Raga-Carbajal E, Olvera C, Rodríguez-Alegría ME, Carrillo-Nava E, Costas M, Munguía AL. 2015. Intrinsic Levanase Activity of Bacillus subtilis 168 Levansucrase (SacB). PLOS ONE 10:e0143394.

49. Cole BJ, Feltcher ME, Waters RJ, Wetmore KM, Mucyn TS, Ryan EM, Wang G, Ul-Hasan S, McDonald M, Yoshikuni Y, Malmstrom RR, Deutschbauer AM, Dangl JL, Visel A. 2017. Genome-wide identification of bacterial plant colonization genes. PLOS Biology 15:e2002860.

50. Liu Z, Beskrovnaya P, Melnyk RA, Hossain SS, Khorasani S, O’Sullivan LR, Wiesmann CL, Bush J, Richard JD, Haney CH. 2018. A Genome-Wide Screen Identifies Genes in Rhizosphere-Associated Pseudomonas Required to Evade Plant Defenses. mBio 9:e00433–18.

51. Budiharjo A, Chowdhury SP, Dietel K, Beator B, Dolgova O, Fan B, Bleiss W, Ziegler J, Schmid M, Hartmann A, Borriss R. 2014. Transposon Mutagenesis of the Plant-Associated Bacillus amyloliquefaciens ssp. plantarum FZB42 Revealed That the nfrA and RBAM17410 Genes Are Involved in Plant-Microbe-Interactions. PLoS One 9:e98267.

52. Guerra-Garcia FJ, Sankari S. 2025. A CRISPR Interference System for Inducible Gene Knockdown in soil bacterium Sinorhizobium meliloti. bioRxiv 10.1101/2025.09.23.678109.

53. Bellahsen O, Díaz-Méndez R, Romero D. 2025. Genomic engineering in Rhizobium etli: implementation and evaluation of systems based on dCas9. Front Microbiol 16:1604430.

54. Gao S, Wu H, Yu X, Qian L, Gao X. 2016. Swarming motility plays the major role in migration during tomato root colonization by *Bacillus subtilis* SWR01. Biological Control 98:11–17.

55. Dietel K, Beator B, Budiharjo A, Fan B, Borriss R. 2013. Bacterial Traits Involved in Colonization of Arabidopsis thaliana Roots by Bacillus amyloliquefaciens FZB42. Plant Pathol J 29:59–66.

56. Nordgaard M, Blake C, Maróti G, Hu G, Wang Y, Strube ML, Kovács ÁT. 2022. Experimental evolution of Bacillus subtilis on Arabidopsis thaliana roots reveals fast adaptation and improved root colonization. iScience 25:104406.

57. Bucher T, Oppenheimer-Shaanan Y, Savidor A, Bloom-Ackermann Z, Kolodkin-Gal I. 2015. Disturbance of the bacterial cell wall specifically interferes with biofilm formation. Environ Microbiol Rep 7:990–1004.

58. Brown S, Santa Maria JP, Walker S. 2013. Wall Teichoic Acids of Gram-Positive Bacteria. Annu Rev Microbiol 67:10.1146/annurev-micro-092412–155620.

59. Xu Z, Zhang H, Sun X, Liu Y, Yan W, Xun W, Shen Q, Zhang R. 2019. Bacillus velezensis Wall Teichoic Acids Are Required for Biofilm Formation and Root Colonization. Appl Environ Microbiol 85:e02116–18.

60. Bhattacharyya A, Pablo CHD, Mavrodi OV, Weller DM, Thomashow LS, Mavrodi DV. 2021. Rhizosphere plant-microbe interactions under water stress. Adv Appl Microbiol 115:65–113.

61. Miller KJ, Wood JM. 1996. Osmoadaptation by rhizosphere bacteria. Annu Rev Microbiol 50:101–136.

62. Chuberre C, Plancot B, Driouich A, Moore JP, Bardor M, Gügi B, Vicré M. 2018. Plant Immunity Is Compartmentalized and Specialized in Roots. Front Plant Sci 9:1692.

63. Hashem A, Tabassum B, Fathi Abd_Allah E. 2019. Bacillus subtilis: A plant-growth promoting rhizobacterium that also impacts biotic stress. Saudi J Biol Sci 26:1291–1297.

64. Gilhar O, Ben-Navi LR, Olender T, Aharoni A, Friedman J, Kolodkin-Gal I. 2024. Multigenerational inheritance drives symbiotic interactions of the bacterium *Bacillus subtilis* with its plant host. Microbiological Research 286:127814.

65. Tsai H-H, Wang J, Geldner N, Zhou F. 2023. Spatiotemporal control of root immune responses during microbial colonization. Current Opinion in Plant Biology 74:102369.

66. Bhavsar AP, Erdman LK, Schertzer JW, Brown ED. 2004. Teichoic Acid Is an Essential Polymer in Bacillus subtilis That Is Functionally Distinct from Teichuronic Acid. J Bacteriol 186:7865–7873.

67. Roney IJ, Rudner DZ. 2024. Bacillus subtilis uses the SigM signaling pathway to prioritize the use of its lipid carrier for cell wall synthesis. PLOS Biology 22:e3002589.

68. Wang B, Teng Z, Siersma T, Kloet F van der, Hamoen L. 2026. Transposon library anomaly reveals importance of cell wall teichoic acids for kin discrimination. bioRxiv 10.64898/2026.07.07.736938.

69. Lacroix EM, Aeppli M, Boye K, Brodie E, Fendorf S, Keiluweit M, Naughton HR, Noël V, Sihi D. 2023. Consider the Anoxic Microsite: Acknowledging and Appreciating Spatiotemporal Redox Heterogeneity in Soils and Sediments. ACS Earth Space Chem 7:1592–1609.

70. Højberg O, Schnider U, Winteler HV, Sørensen J, Haas D. 1999. Oxygen-Sensing Reporter Strain of Pseudomonas fluorescens for Monitoring the Distribution of Low-Oxygen Habitats in Soil. Appl Environ Microbiol 65:4085–4093.

71. Fang FC. 1997. Perspectives series: host/pathogen interactions. Mechanisms of nitric oxide-related antimicrobial activity. J Clin Invest 99:2818–2825.

72. Zemojtel T, Fröhlich A, Palmieri MC, Kolanczyk M, Mikula I, Wyrwicz LS, Wanker EE, Mundlos S, Vingron M, Martasek P, Durner J. 2006. Plant nitric oxide synthase: a never-ending story? Trends in Plant Science 11:524–525.

73. López-Gómez P, Buezo J, Urra M, Cornejo A, Esteban R, Fernández de los Reyes J, Urarte E, Rodríguez-Dobreva E, Chamizo-Ampudia A, Eguaras A, Wolf S, Marino D, Martínez-Merino V, Moran JF. 2024. A new oxidative pathway of nitric oxide production from oximes in plants. Molecular Plant 17:178–198.

74. Ordal GW, Villani DP, Rosendahl MS. 1979. Chemotaxis Towards Sugars by Bacillus subtilis. Microbiology 115:167–172.

75. Hafner BD, Pietz O, King WL, Scharfetter JB, Bauerle TL. 2025. Early developmental shifts in root exudation profiles of five *Zea mays* L. genotypes. Plant Science 354:112439.

76. Hemelda NM, Noutoshi Y. 2025. Root-exuded sugars as drivers of rhizosphere microbiome assembly. Plant Biotechnology 42:215–227.

77. Carvalhais LC, Dennis PG, Badri DV, Kidd BN, Vivanco JM, Schenk PM. 2015. Linking Jasmonic Acid Signaling, Root Exudates, and Rhizosphere Microbiomes. Mol Plant Microbe Interact 28:1049–1058.

78. Loo EP-I, Durán P, Pang TY, Westhoff P, Deng C, Durán C, Lercher M, Garrido-Oter R, Frommer WB. 2024. Sugar transporters spatially organize microbiota colonization along the longitudinal root axis of *Arabidopsis*. Cell Host & Microbe 32:543–556.e6.

79. Guo W-J, Nagy R, Chen H-Y, Pfrunder S, Yu Y-C, Santelia D, Frommer WB, Martinoia E. 2014. SWEET17, a Facilitative Transporter, Mediates Fructose Transport across the Tonoplast of Arabidopsis Roots and Leaves. Plant Physiol 164:777–789.

80. Chen H-Y, Huh J-H, Yu Y-C, Ho L-H, Chen L-Q, Tholl D, Frommer WB, Guo W-J. 2015. The Arabidopsis vacuolar sugar transporter SWEET2 limits carbon sequestration from roots and restricts Pythium infection. The Plant Journal 83:1046–1058.

81. Chardon F, Bedu M, Calenge F, Klemens PAW, Spinner L, Clement G, Chietera G, Léran S, Ferrand M, Lacombe B, Loudet O, Dinant S, Bellini C, Neuhaus HE, Daniel-Vedele F, Krapp A. 2013. Leaf Fructose Content Is Controlled by the Vacuolar Transporter SWEET17 in *Arabidopsis*. Current Biology 23:697–702.

82. Boonstra M, de Jong IG, Scholefield G, Murray H, Kuipers OP, Veening J-W. 2013. Spo0A regulates chromosome copy number during sporulation by directly binding to the origin of replication in Bacillus subtilis. Molecular Microbiology 87:925–938.

83. Koo B-M, Kritikos G, Farelli JD, Todor H, Tong K, Kimsey H, Wapinski I, Galardini M, Cabal A, Peters JM, Hachmann A-B, Rudner DZ, Allen KN, Typas A, Gross CA. 2017. Construction and Analysis of Two Genome-Scale Deletion Libraries for Bacillus subtilis. Cell Syst 4:291–305.e7.

84. Sewgoolam B, Janssen AB, Martin LS, Rengifo-Gonzalez M, Bakker V de, Rozendal B, Cremers AJH, Veening J-W. 2026. A functional genetic landscape of antibiotic sensitivity across the pneumococcal pangenome reveals conserved and lineage-specific vulnerabilities. bioRxiv 10.64898/2026.02.26.708248.

85. Jaiaue P, Srimongkol P, Thitiprasert S, Tanasupawat S, Cheirsilp B, Assabumrungrat S, Thongchul N. 2021. A modified approach for high-quality RNA extraction of spore-forming Bacillus subtilis at varied physiological stages. Mol Biol Rep 48:6757–6768.

