## Supplementary File for "Genome-wide interrogation of genetic requirements for root colonization by CRISPRi-seq in *Bacillus subtilis*"

Table S1. Bacterial strains used in this study.

| **Species** | **Strain** | **Source** |
| --- | --- | --- |
| *B. subtilis* | *Bacillus subtilis* 168, trhC::pspank-dcas9 | This study |
| *B. subtilis* | 168, thrC::pSpank-dCas9, *amyE*::P3-sgRNA_library | This study |
| *B. subtilis* | NCIB 3610 ComIQ12L, thrC::pSpank-dCas9, *amyE*::P3-sgRNA_library | This study |
| *B. subtilis* | NCIB 3610 ComIQ12L, *amyE*::PrrnB-GFP-Cm | This study |
| *B. subtilis* | NCIB 3610 ComIQ12L, *lacA::*PrrnB::mScarlet-I3 | This study |
| *B. subtilis* | NCIB 3610 ComIQ12L, thrC::pspank-dcas9, amyE::p3-sgRNA murF | This study |
| *B. subtilis* | NCIB 3610 ComIQ12L, thrC::pspank-dcas9, amyE::p3-sgRNA ganA | This study |
| *B. subtilis* | NCIB 3610 ComIQ12L, thrC::pspank-dcas9, amyE::p3-sgRNA dnaA | This study |
| *B. subtilis* | NCIB 3610 ComIQ12L, Δ*yopQ*-Kan | This study |
| *B. subtilis* | NCIB 3610 ComIQ12L, Δ*lex*A-Kan | This study |
| *B. subtilis* | NCIB 3610 ComIQ12L, Δ*cdaA*-Kan | This study |
| *B. subtilis* | NCIB 3610 ComIQ12L, Δ*tagE*-Kan | This study |
| *B. subtilis* | NCIB 3610 ComIQ12L, Δ*hag*-Kan | This study |
| *B. subtilis* | NCIB 3610 ComIQ12L, *ΔreoM*-Kan | This study |
| *B. subtilis* | NCIB 3610 ComIQ12L, Δ*ecsB*-Kan | This study |
| *B. subtilis* | NCIB 3610 ComIQ12L, Δ*hemAT*-Kan | This study |
| *B. subtilis* | NCIB 3610 ComIQ12L, Δ*rghR*-Kan | This study |
| *B. subtilis* | NCIB 3610 ComIQ12L, Δ*cheC*-Kan | This study |
| *B. subtilis* | NCIB 3610 ComIQ12L, Δ*dltA-*Kan | This study |
| *B. subtilis* | NCIB 3610 ComIQ12L, *amyE*::PrrnB-GFP-Cm, Δ*yopQ*-Kan | This study |
| *B. subtilis* | NCIB 3610 ComIQ12L, *amyE*::PrrnB-GFP-Cm, Δ*lex*A-Kan | This study |
| *B. subtilis* | NCIB 3610 ComIQ12L, *amyE*::PrrnB-GFP-Cm, Δ*cdaA*-Kan | This study |
| *B. subtilis* | NCIB 3610 ComIQ12L, *amyE*::PrrnB-GFP-Cm, Δ*tagE*-Kan | This study |
| *B. subtilis* | NCIB 3610 ComIQ12L, *amyE*::PrrnB-GFP-Cm, Δ*hag*-Kan | This study |
| *B. subtilis* | NCIB 3610 ComIQ12L, *amyE*::PrrnB-GFP-Cm, *ΔreoM*-Kan | This study |
| *B. subtilis* | NCIB 3610 ComIQ12L, *amyE*::PrrnB-GFP-Cm, Δ*ecsB*-Kan | This study |
| *B. subtilis* | NCIB 3610 ComIQ12L, *amyE*::PrrnB-GFP-Cm, Δ*hemAT*-Kan | This study |
| *B. subtilis* | NCIB 3610 ComIQ12L, *amyE*::PrrnB-GFP-Cm, Δ*rghR*-Kan | This study |
| *B. subtilis* | NCIB 3610 ComIQ12L, *amyE*::PrrnB-GFP-Cm, Δ*cheC*-Kan | This study |
| *B. subtilis* | NCIB 3610 ComIQ12L, *amyE*::PrrnB-GFP-Cm, Δ*dltA-*Kan | This study |

Table S2. Primers used in this study.

| **Primer** | **Sequence** | **Source** |
| --- | --- | --- |
| OVL6453 | CGAAAGCACCTGCTGGAagggatcctagaagcttatcgaattct | This study |
| OVL6454 | CGAAAGCACCTGCTGGAttactagtaaaatctaagacatatcatgat | This study |
| OVL6455 | CGAAAGCACCTGCCCATgtaattcggtcgacagatcttcgtcgg | This study |
| OVL6456 | CGAAAGCACCTGCCCATccctcgagcaagcagaagacggcatac | This study |
| OVL6490 | CCGGCGCAGAAGTTTGAACGAAAAG | Sewgoolam et al., 2026 |
| OVL6491 | AGGAGCCCGAATCCATCCTCGAATA | Sewgoolam et al., 2026 |
| OVL6528 | AGTAGCGGTCTCCAAGACAGCTAGCCGCATGCAAGCTAAT | This study |
| OVL6529 | AGTAGCGGTCTCCTTCCTCCTGCTTAATTGTTATCCGCTCACAATT | This study |
| OVL6530 | AGTAGCGGTCTCGGGAAAAAATGGATAAGAAATACTCAATAGGCT | This study |
| OVL6531 | AGTAGCGGTCTCGTCTTAGTCACCTCCTAGCTGACTCAAA | This study |
| OVL6581 | AATTCATGTAAAAGATGAGGTTGG | This study |
| OVL6582 | TCATACACGGGCCGCTCCTTTTAC | This study |
| OVL6759 | aataacgtctcggtttaagagctatgctgg | This study |
| OVL6760 | ctcctcgtctcatatagttattataccaggggg | This study |
| oVM_019_yopQ_FF | gatgccttcatcaactagaagg | This study |
| oVM_020_yopQ_FR | ggtcttggtctcctccttag | This study |
| oVM_021_lexA_FF | actcttccacaatgctgacg | This study |
| oVM_022_lexA_FR | gtcgtatatgtggatgatcagc | This study |
| oVM_023_ybbP_FF | atcgtcgcagatggtgtt | This study |
| oVM_024_ybbP_FR | cctgatgtattgctgctgtta | This study |
| oVM_025_tagE_FF | gctcctccttcctttaagattct | This study |
| oVM_026_tagE_FR | caggctatagtcgtttactctg | This study |
| oVM_027_yrzL_FF | tcttggaactcatgtcaatcag | This study |
| oVM_028_yrzl_FR | tatagctccggctttggttt | This study |
| oVM_029_rghRA_FF | catgtcacgctgtatcagac | This study |
| oVM_030_rghRA_FR | gtgtggatgtactatatatcggc | This study |
| oVM_031_hag_FF | gtcacagcttgatgtgcag | This study |
| oVM_032_hag_FR | gctgatactcctagatctgagc | This study |
| oVM_033_ecsB_FF | cgattcctgcgataaacacc | This study |
| oVM_034_ecsB_FR | aaccatcctgcagaaacga | This study |
| oVM_035_hemAT_FF | agccgctttatcatatgctg | This study |
| oVM_036_hemAT_FR | tctgtacaggtaaagctctcc | This study |
| oVM_039_cheC_FF | tatcacagcggcaaccatg | This study |
| oVM_040_cheC_FR | ctgtgcaatcagttcatcct | This study |
| oVM_041_dltA_FF | catgtactagacgtccctgtc | This study |
| oVM_042_dltA_FR | tgttgctgactctttcacac | This study |

Table S3. Plasmids used in this study.

| **Plasmid** | **Relevant characteristics** | **Source** |
| --- | --- | --- |
| pMB002 | a pMB001 derivative with a MCS instead kinA; bla, thrC, Pspank, erm | Boonstra *et al*., 2013 |
| pJWV102-PL-dCas9 | Plac‐dcas9sp. PczcD-GFP, tetM | Liu et al., 2017 |
| pDS22 | read1-P3-mCherry-dCas9 handle-read2-P7; spc | This study |

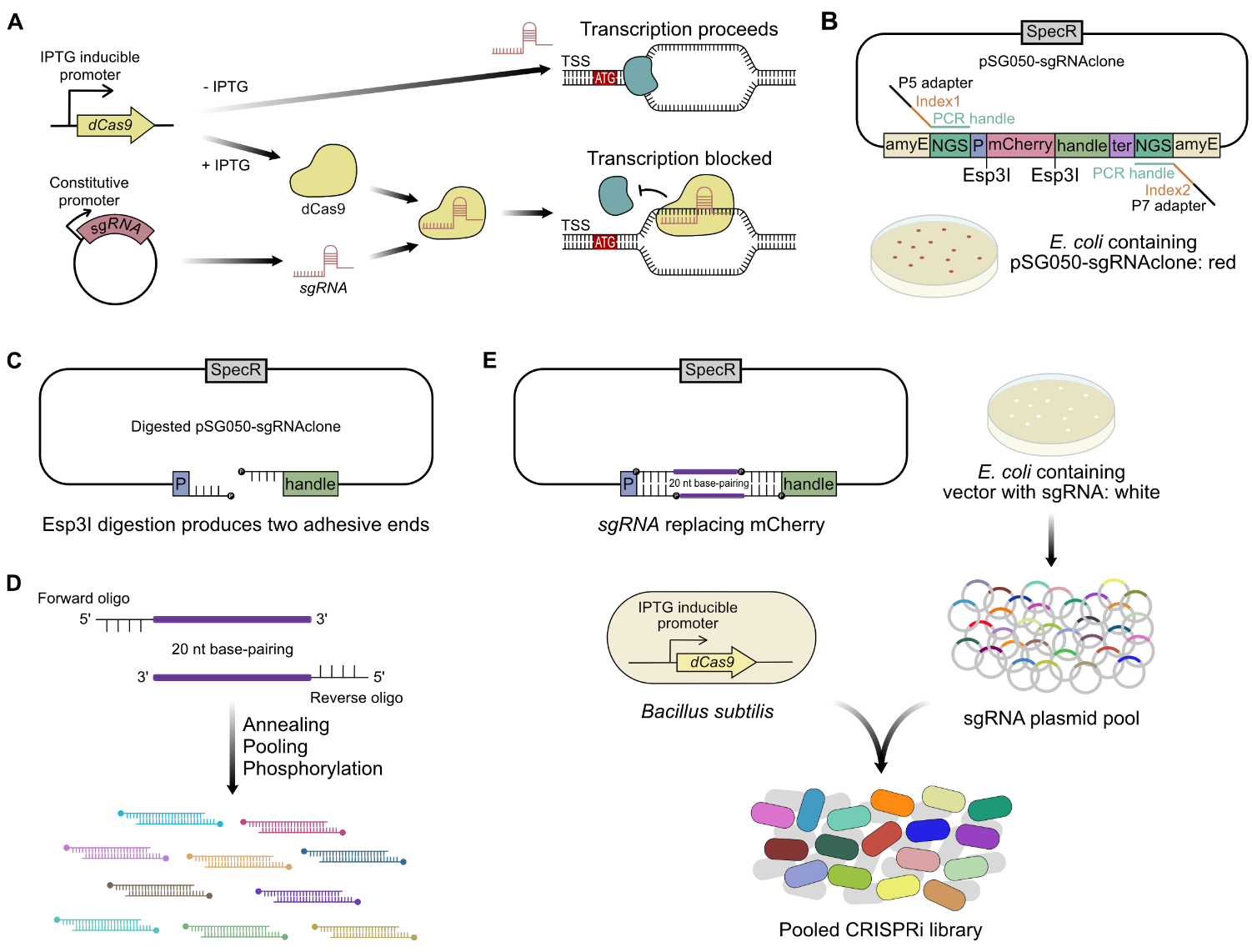

**Figure S1. Schematic of *B. subtilis* CRISPRi-seq library construction and plant colonization workflow.** **(A) CRISPRi mechanism and design strategy.** An IPTG-inducible *dCas9* and a constitutively expressed sgRNA targeting the region downstream of the transcriptional start site (TSS) are used to block transcript elongation. **(B) Map of the cloning vector pSG050-sgRNAclone.** The vector features *amyE* homology arms for genomic integration, a SpecR resistance marker, and an mCherry cassette flanked by Esp3I restriction sites for Golden Gate-based cloning of sgRNA spacers. **(C) Vector preparation.** Esp3I digestion removes the mCherry cassette, producing two distinct adhesive ends for directional ligation. **(D) sgRNA pool generation.** Forward and reverse oligonucleotides containing 20 nt base-pairing sequences that target each gene/operon of the *B. subtilis genomes*, containing Esp3I overhangs, are annealed, phosphorylated, and pooled at equimolar ratios. **(E) Library cloning and selection.** The sgRNA pool is ligated into the digested vector and transformed into *E. coli*. Successful insertion replaces mCherry, allowing for a red-to-white colony screen. The resulting plasmid library is purified and transformed into *B. subtilis* 3610 containing the IPTG inducible dCas9, where the CRISPRi construct integrates into the *amyE* locus to generate the pooled CRISPRi library.

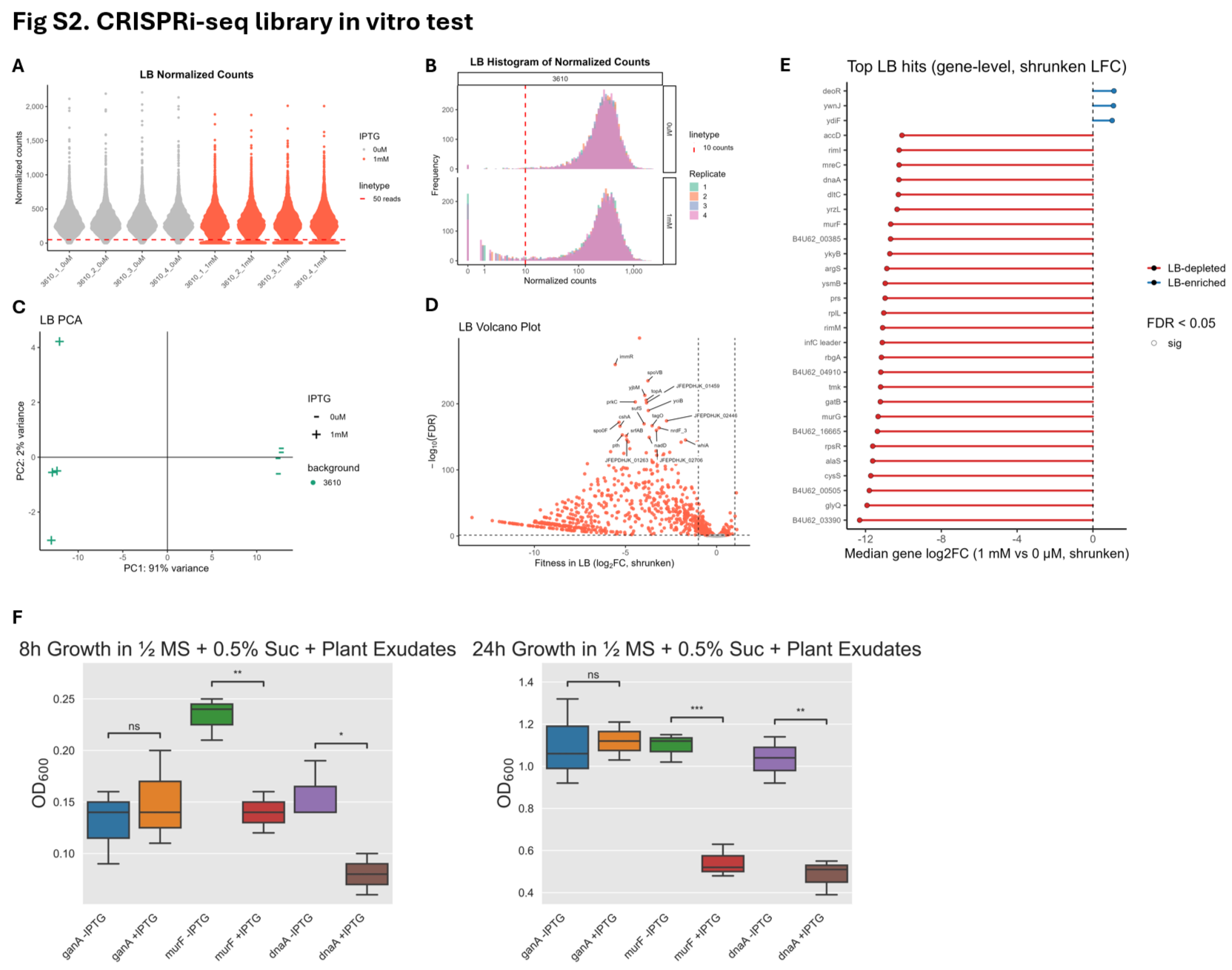

**Figure S2. Quality control and analysis of the 3610 CRISPRi-seq library grown in vitro in LB.** The library was cultured for approximately 12 generations in LB medium in the presence (1 mM) or absence (0 mM) of IPTG prior to sequencing. **(A)** Violin plot comparing normalized sgRNA read counts in induced (1 mM IPTG) versus uninduced (0 µM IPTG) conditions. Each dot represents a single sgRNA. **(B)** Histogram showing the distribution of normalized sgRNA read counts. The x-axis represents read counts, and the y-axis indicates the frequency (number of sgRNAs). **(C)** Principal Component Analysis (PCA) of the of rlog normalized Deseq data set. Shapes indicate treatment condition (+: 1 mM IPTG; −: 0 mM IPTG). **(D)** Volcano plot of DESeq2 analysis comparing 1 mM vs 0 mM IPTG. Orange dots represent genes meeting the significance threshold (Log2FC < -1 and FDR < 0.05). **(E)** Lollipop plot displaying the top-ranking gene targets identified in the screen. Color coding indicates the effect on fitness (Blue: Costly; Red: Essential). **(F)** Validation of dCas9 knockdown with individual sgRNAs. Optical density at 600 nm of B. subtilis 3610 dCas9 strains carrying individual sgRNAs against the essential genes murF and dnaA or the non-essential gene ganA, measured 8 h and 24 h after inoculation at a starting OD₆₀₀ of 0.05 in plant exudate medium supplemented with 15 mM sucrose, with or without 1 mM IPTG. Boxes show the median and interquartile range. n = 3 biological replicates per strain. Each mutant was compared with the wild type by exact two-sided Mann-Whitney U test, with p-values adjusted within the panel by the Benjamini-Hochberg procedure (** adjusted p < 0.01).

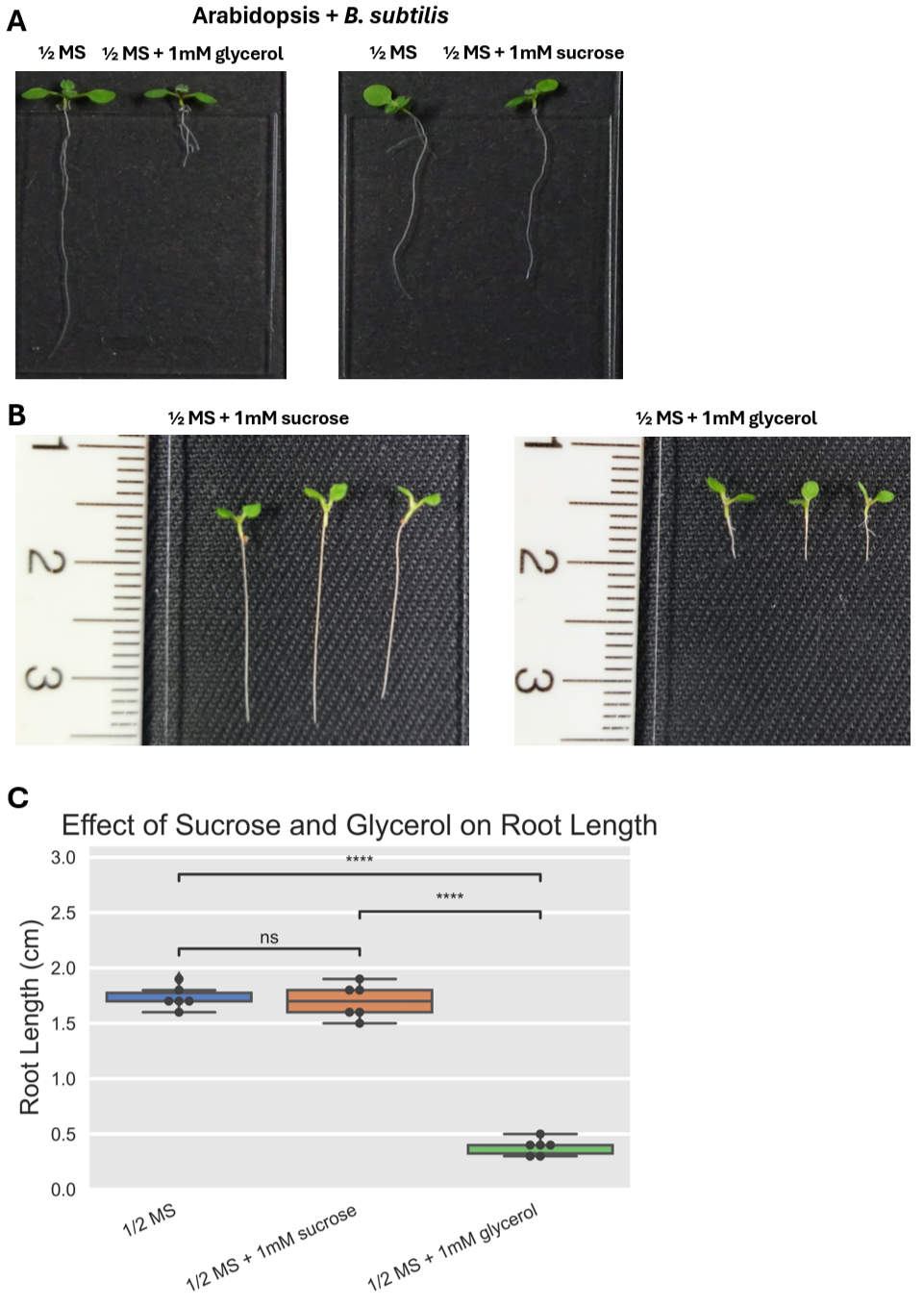

**Figure S3. Impact of glycerol and sucrose supplementation on *Arabidopsis* root elongation.** Seedlings were germinated on ½ MS agar for 3 days and transferred to liquid ½ MS medium supplemented with 1 mM glycerol or 1 mM sucrose for an additional 3 days. **(A, B)** Representative images of root phenotypes after 6 days of total growth. **(C)** Quantification of primary root length. Data are presented as mean ± SD (n = 5 biological replicates). Statistical significance was determined by independent t-test.

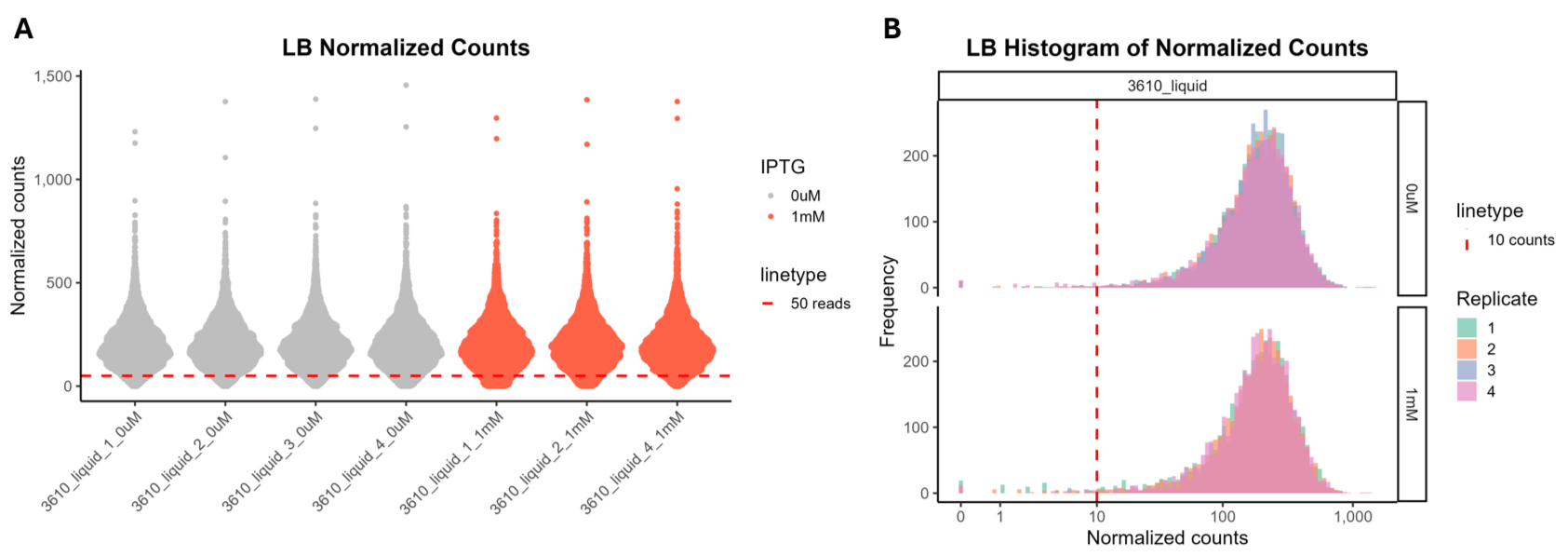

**Figure S4. Control of 3610 CRISPRi-seq library following pre-induction.** TThe library was cultured in LB medium for approximately 4 generations in the presence (1 mM) or absence (0 mM) of IPTG to pre-induce the CRISPRi system prior to hydroponic inoculation. **(A)** Violin plot comparing normalized sgRNA read counts in pre-induced (1 mM IPTG) versus uninduced (0 µM IPTG) conditions. Each dot represents a single sgRNA. **(B)** Histogram showing the distribution of normalized sgRNA read counts. The x-axis represents read counts, and the y-axis indicates the frequency (number of sgRNAs).

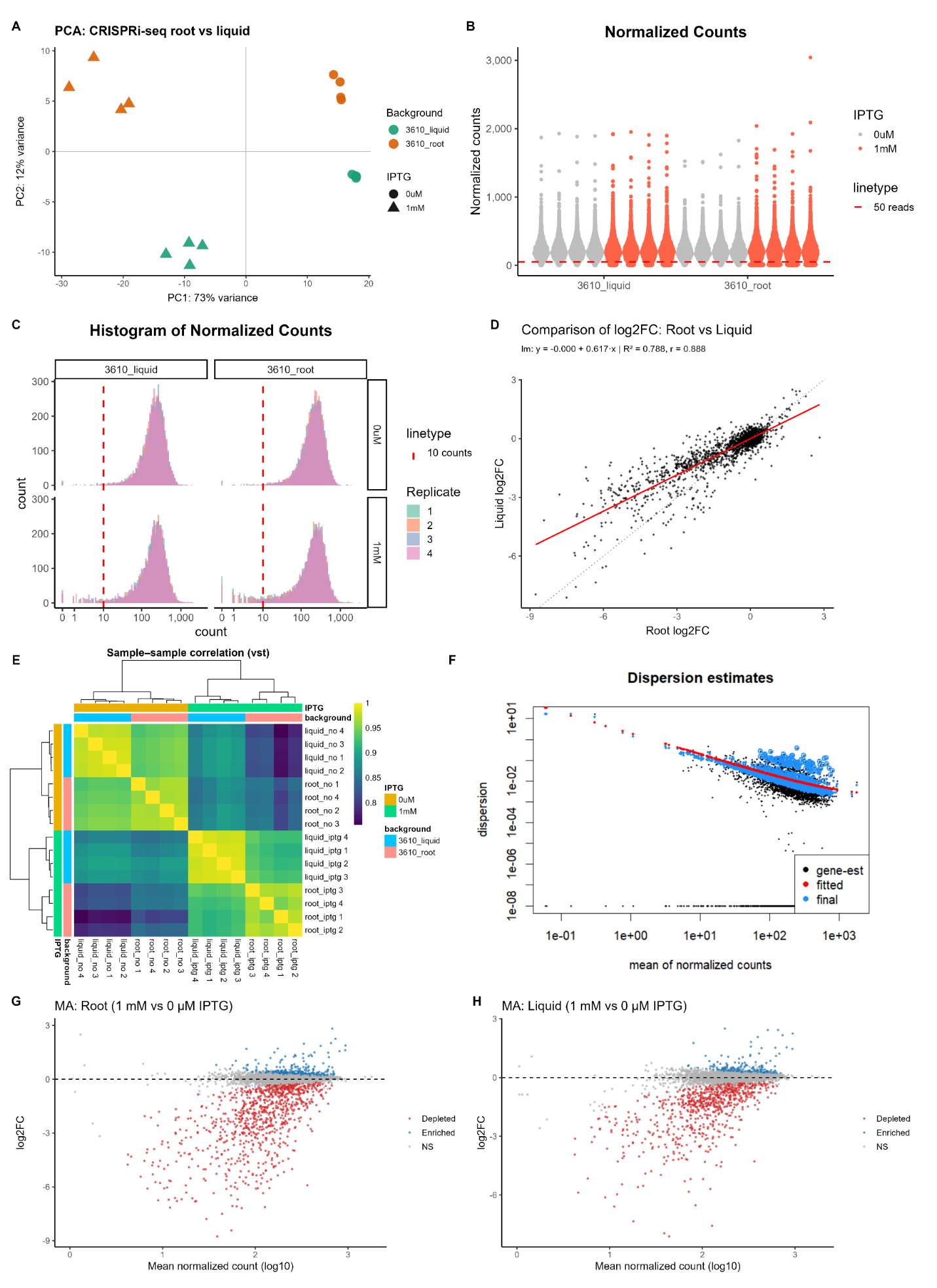

**Figure S5. Quality control of root-colonized versus non-colonized CRISPRi-seq libraries.** Comparisons were made between libraries that were cultured in the hydroponics system that either colonized *Arabidopsis* roots or did not colonize and were in the surrounding liquid, in the presence (1 mM) or absence (0 mM) of IPTG. **(A)** Violin plot comparing normalized sgRNA read counts of uncolonized (3610_liquid) and root colonized (3610_root) samples in both induced 1 mM IPTG (orange) versus uninduced 0 mM IPTG (grey) conditions. Each dot represents a single sgRNA. **(B)** Histogram showing the distribution of normalized sgRNA read counts. The x-axis represents read counts, and the y-axis indicates the frequency (number of sgRNAs) for 3610_root and 3610_liquid conditions **(C)**. Scatter plot comparing the Log2 Fold Change (|log_2_FC|) of sgRNAs in root samples (x-axis) versus liquid samples (y-axis). The red solid line represents the linear regression fit. The regression equation, coefficient of determination (R^2^), and Pearson correlation coefficient (r) are displayed in the top left. The dotted line represents the identity line (x=y). **(D)** Sample-to-sample correlation heatmap based on variance-stabilizing transformed (VST) data. The color scale indicates correlation strength (yellow = high; blue = low). Annotation bars on the right and top denote the experimental condition (IPTG) and sample origin (Background: Liquid vs Root). **(E)** Dispersion plot showing the relationship between mean normalized counts (x-axis) and dispersion (y-axis). This visualizes the shrinkage of gene-wise dispersion estimates (black dots) toward the fitted trend line (red line) to generate the final estimates (blue dots) used for statistical testing. **(F)** MA plots displaying differential sgRNA abundance (1 mM vs 0 mM IPTG) for Root (left) and Liquid (right) conditions. The y-axis represents (|log_2_FC|) and the x-axis represents mean normalized counts. Red dots indicate significantly depleted or enriched sgRNAs. **(G)** Heatmap of log-transformed [log10(x+1)] DEseq2 normalized read counts for all sgRNAs. Rows represent individual sgRNAs and columns represent samples. Annotation bars indicate replicate number, IPTG condition, and growth environment.

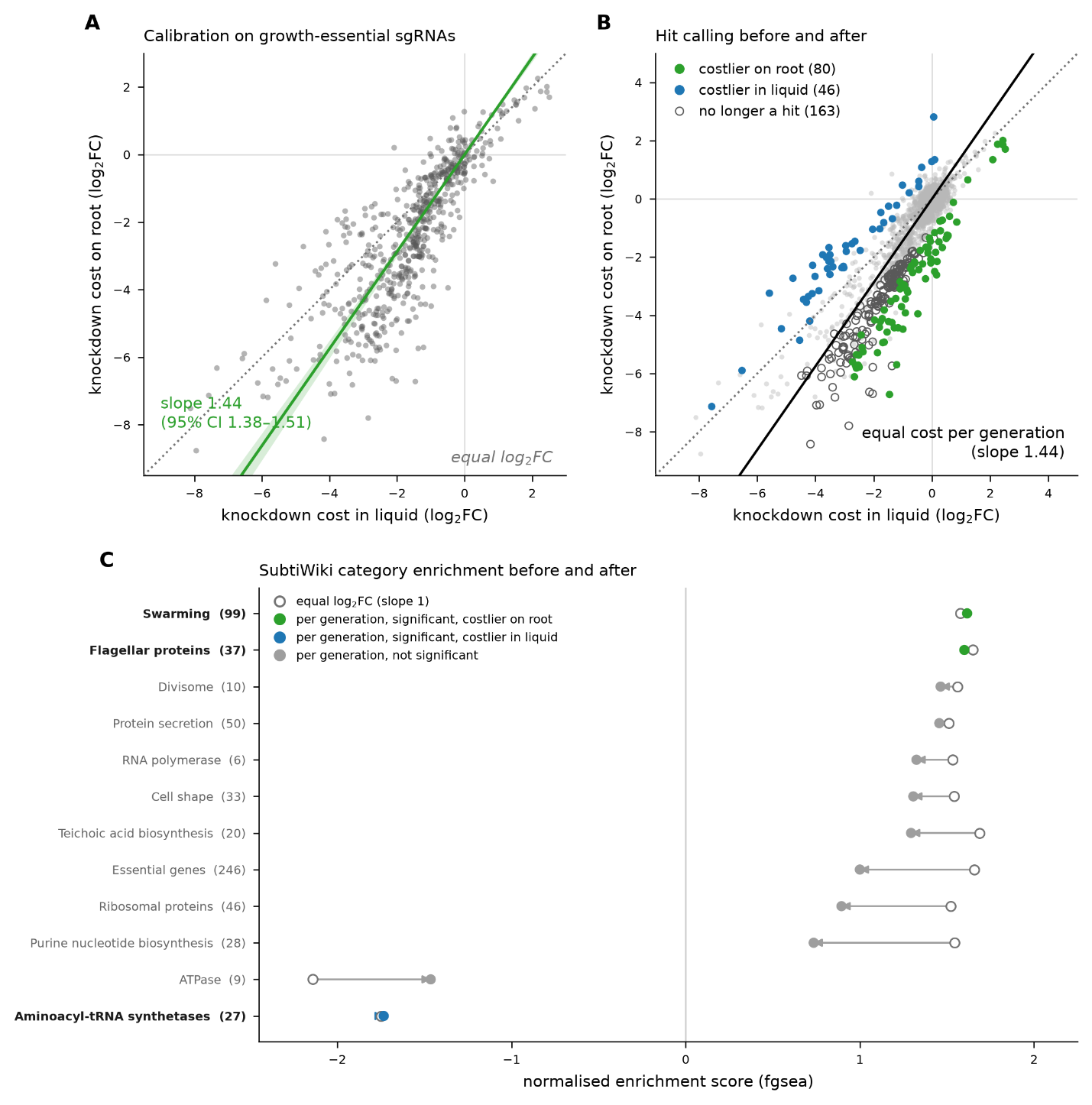
**Figure S6. Knockdown costs compared per generation.** Root-attached and free-swimming populations may go through different numbers of generations during the 24 h experiment. If so, the faster-dividing population would show a larger log₂FC even for knockdowns that are equally costly per generation. To correct for this, we calibrated the root-to-liquid generation ratio on the sgRNAs most strongly costly in the LB screen and repeated the root-versus-liquid comparison per generation (Data S7). **(A)** For the 599 sgRNAs that are significant and costly (|log₂FC| ≤ -1, FDR < 0.05) in the LB screen (Data S1), and whose cost is therefore expected to be dominated by growth rate, the IPTG-induction effect in liquid is plotted against the effect on roots. Green, Deming regression through the origin (band, 95% bootstrap CI); the slope of 1.44 (95% CI 1.38-1.51) estimates the ratio of generations on the root to generations in liquid. Dotted, the identity line used in Figure 2A. **(B)** All 4,326 sgRNAs. Filled points are significant against the calibrated line (FDR < 0.05, |log₂FC| > 1): green, required for colonization (n = 80); blue, costly on roots (n = 46). Open circles, the 163 Figure 2A hits that are not significant per generation. The sgRNAs called required fell from 235 under the identity line to 80, of which 72 were also required under the identity line, while those called costly rose from 14 to 46. The cell wall genes *tagE*, *tagA*, *tagB*, *mnaA* and *walRK* remained significant, whereas the essential wall teichoic acid backbone steps *tagO*, *tagD*, *tagF*, *tagG* and *tagH* did not. **(C)** Normalised enrichment scores for every SubtiWiki category significant under either comparison (set size in parentheses): open circles, ranking by the identity-line statistic as in Figure 2D; filled, by the calibrated statistic; bold labels, significant after calibration. Positive scores indicate enrichment among genes relatively costlier on the root; two category names are abbreviated. Motility remained the dominant signature (swarming, FDR = 9.3 × 10^-4^; flagellar proteins, FDR = 9.2 × 10^-5^). Essential genes, ribosomal proteins and purine nucleotide biosynthesis categories were no longer significant.
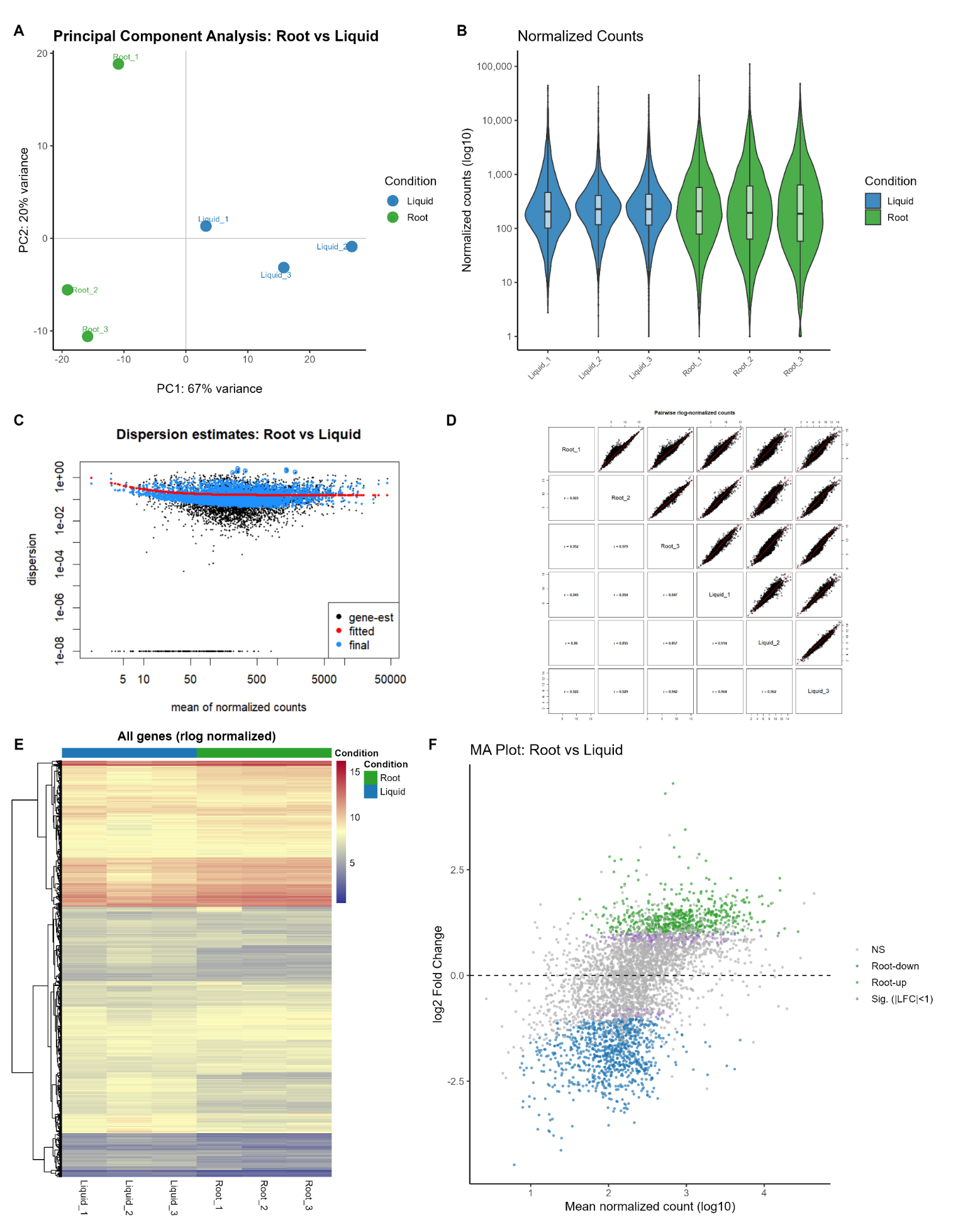
**Figure S7. Quality control of the root-colonized versus non-colonized RNA-seq libraries.** Total RNA was collected from root-attached and free-swimming cells 24 h post-inoculation. Analyses use three biological replicates per condition (Root_1 to Root_3, Liquid_1 to Liquid_3). Root is shown in green and Liquid in blue throughout. **(A)** Principal component analysis of rlog-transformed counts, with samples coloured by condition and labelled by replicate. PC1 accounts for 67% and PC2 for 20% of the variance**. (B)** Violin plots of DESeq2 size-factor-normalized counts per sample, plotted on a log10 scale with axis labels back-transformed to counts, filled by condition, with an embedded box showing the median and interquartile range. Each observation is a single gene. **(C)** DESeq2 dispersion estimates against the mean of normalized counts, showing per-gene estimates (black), the fitted mean-dispersion trend (red) and the final maximum a posteriori estimates used for testing (blue). **(D)** Pairwise comparison of rlog-normalized counts for all six libraries. The diagonal gives the sample name, panels above the diagonal show the scatter of every gene with a locally weighted smoother (red), and panels below the diagonal give the Pearson correlation coefficient for that pair. **(E)** Heatmap of rlog-normalized counts for all genes passing the low-count filter, with genes in rows and samples in columns. Rows are hierarchically clustered; column order is fixed by condition and is not clustered. The colour scale is the rlog value and the annotation bar above the columns denotes condition. **(F)** MA plot of log2 fold change against mean normalized count. Genes are coloured by significance class: root-up (green, padj < 0.05 and log2FC > 1), root-down (blue, padj < 0.05 and log2FC < −1), significant but below the fold-change threshold (purple, padj < 0.05 and absolute log2FC < 1), and not significant (grey).

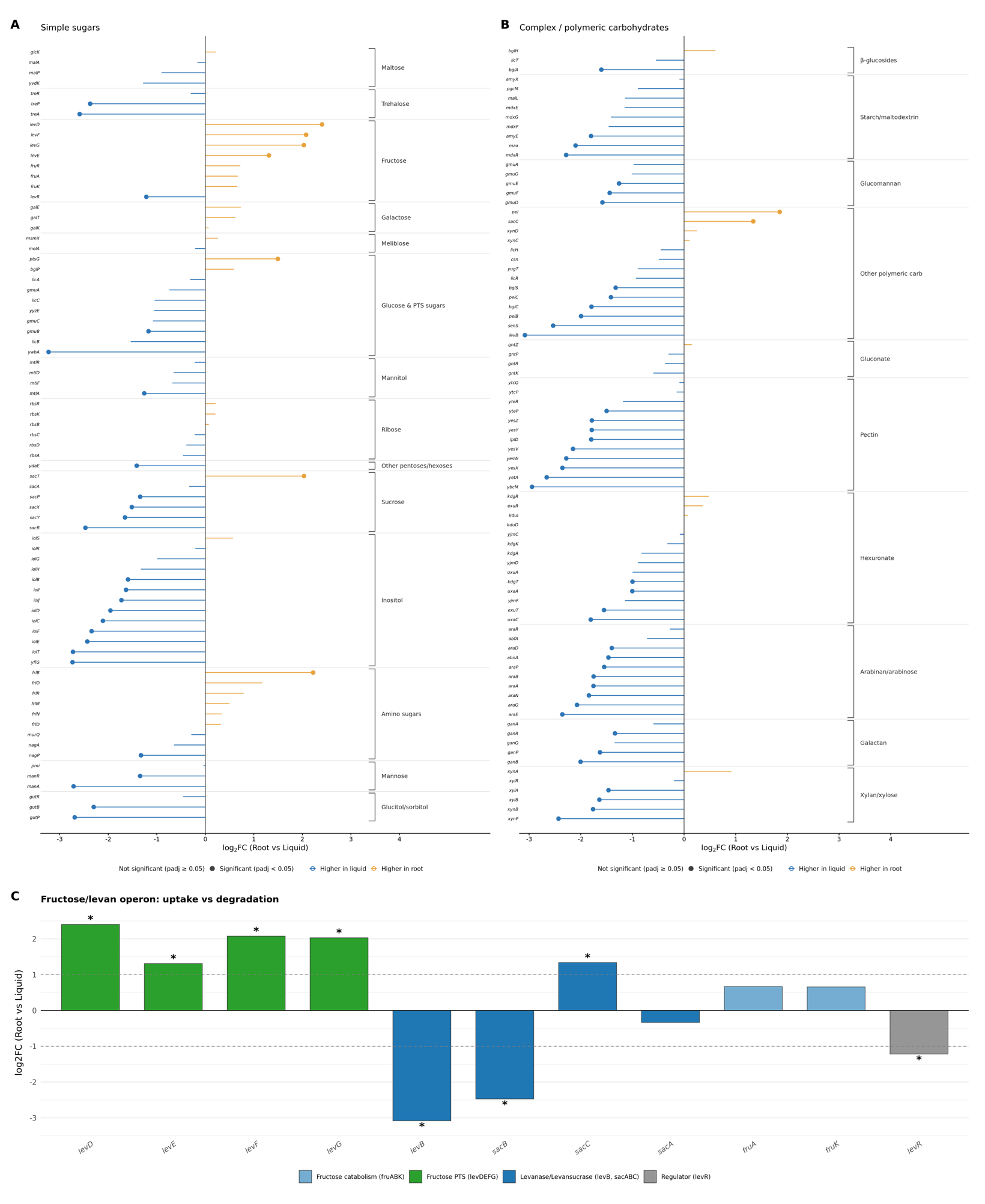

**Figure S8.** Lollipop plots showing the log₂ fold-change (root vs liquid) of *B. subtilis* carbohydrate-utilization genes during hydroponic root colonization (1 dpi). Each point is one gene; the stem runs from zero to its log₂FC. Point colour encodes direction, orange, higher expression during root colonization; blue, higher in the liquid condition, and fill encodes significance: filled points are significant (adjusted *p* < 0.05), hollow points are not. **(A)** Genes for simple-sugar (mono- and disaccharide) utilization, grouped by SubtiWiki sugar category (bracket labels: Maltose, Trehalose, Fructose, Galactose, Melibiose, Glucose & general PTS sugars, Mannitol, Ribose, Other pentoses/hexoses, Sucrose, Inositol, Amino sugars, Mannose, and Glucitol/sorbitol). **(B)** Genes for complex/polymeric carbohydrate utilization, grouped by target substrate (β-glucosides, Starch/maltodextrin, Glucomannan, Other polymeric carbohydrates, Gluconate, Pectin, Hexuronate, Arabinan/arabinose, Galactan, and Xylan/xylose). Genes that SubtiWiki assigns to both a simple and a complex category (e.g. the *lev* fructose/levan operon) are shown only under their simple-sugar category in panel A, so the two panels are non-overlapping. **C)** Bar plot showing the Log2 Fold Change (root vs liquid) of genes belonging to the fructose/levan utilization locus. Genes are grouped by functional role: Fructose PTS transport (*levDEFG*, green), Levanase/Levansucrase (*levB*, *sacABC*, dark blue), Fructose catabolism (*fruABK*, light blue), and the locus regulator (*levR*, grey). Asterisks denote statistical significance (* p-value < 0.05).

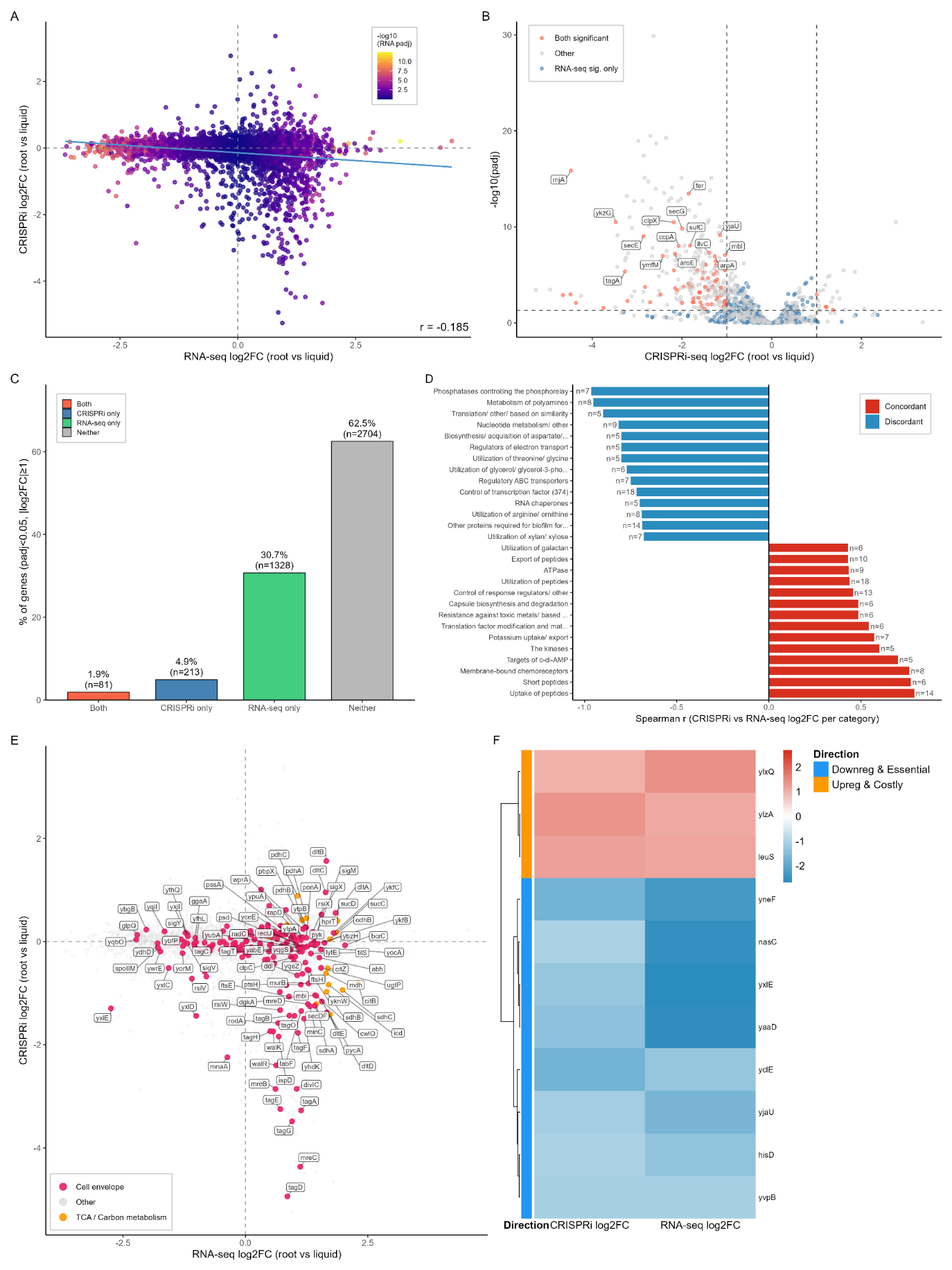

**Figure S9. Extended comparison of CRISPRi-seq fitness effects and RNA-seq differential expression during root colonization.** All panels compare root-colonizing to liquid conditions, as in the main figure. **A)** Genome-wide relationship between CRISPRi-seq and RNA-seq log2FC. Points colored by RNA-seq significance (-log10 padj); blue line shows linear fit. Pearson's r for the genome-wide relationship = -0.185. **B)** CRISPRi-seq volcano plot highlighting RNA-seq-significant genes. Genes are "Both significant" (FDR < 0.05, |log2FC| ≥ 1 in both screens; red), "RNA-seq sig. only" (blue), or "Other" (grey). Dashed lines mark CRISPRi-seq significance and effect-size thresholds. Labels show the top 15 "Both significant" genes, ranked by combined CRISPRi-seq significance and effect size. **C)** Proportional overlap of significant hits. Percentage of genes meeting significance thresholds (FDR < 0.05, |log2FC| ≥ 1) in the CRISPRi-seq screen, RNA-seq screen, both, or neither, with gene counts (n) annotated. **D)** Per-category concordance between CRISPRi-seq and RNA-seq log2FC. Spearman r between CRISPRi-seq and RNA-seq log2FC computed per SubtiWiki category (≥5 genes); the 15 most concordant and 15 most discordant categories are shown, annotated with gene counts (n). Some category names are truncated for display. **E)** Focused comparison of cell envelope and carbon metabolism genes, excluding motility genes. CRISPRi-seq log2FC vs. RNA-seq log2FC for cell envelope (pink) and TCA/carbon metabolism (orange) genes; other genes shown in grey. Labels restricted to genes in these categories meeting significance thresholds. **F)** Heatmap of genes with same-direction significant changes in both screens. Log2FC values for genes meeting significance thresholds (FDR < 0.05, |log2FC| ≥ 1) in both screens with concordant sign, split into "Downreg & Essential" and "Upreg & Costly" groups (row annotation), clustered by log2FC similarity within each group. Complements the opposite-direction convergent set shown in the main figure.

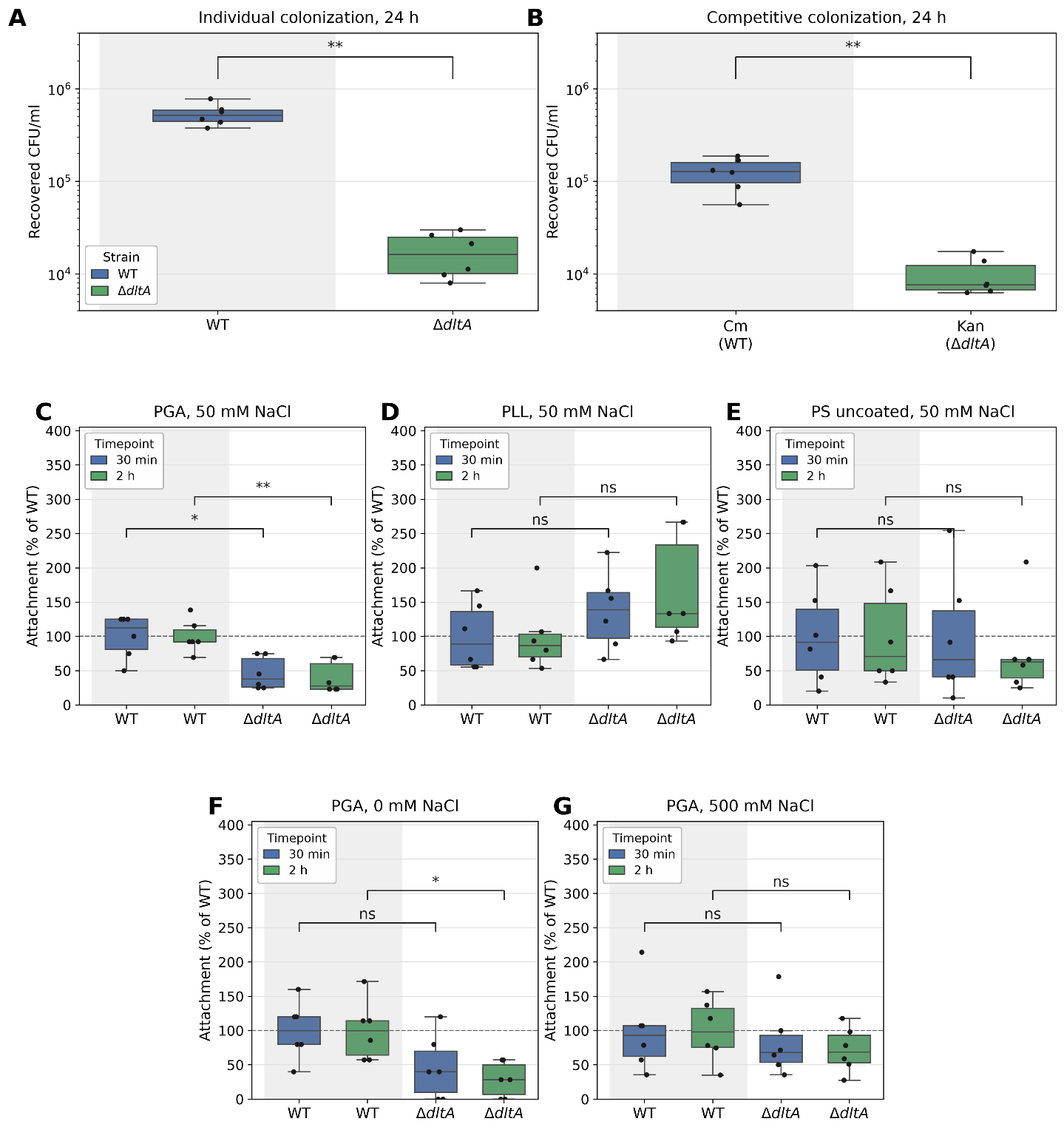

**Figure S10. ΔdltA root colonization, surface specificity and ionic-strength dependence of the attachment defect. (A)** Individual root colonization. Wild type and ΔdltA were inoculated separately into the hydroponic system at OD₆₀₀ 0.02 and roots harvested at 24 h. **(B)** Competitive root colonization. Wild type (Cmʳ) and ΔdltA (Kanʳ) were co-inoculated at OD₆₀₀ 0.01 per strain and both populations enumerated in parallel from the same root homogenate at 24 h. **(C-G)** Attachment of wild type and ΔdltA to charge-defined inert surfaces at 30 min (blue) and 2 h (green), expressed as a percentage of the mean wild-type value on the same surface, in the same buffer and at the same timepoint. **(C)** PGA-coated wells, a negatively charged surface, in base assay buffer; the same data are shown in Figure 6B. **(D)** Poly-L-lysine-coated wells, a positively charged surface, in base assay buffer. **(E)** Uncoated tissue-culture polystyrene, a moderately negative reference surface, in base assay buffer. **(F)** PGA-coated wells with no added NaCl. **(G)** PGA-coated wells in buffer containing 500 mM NaCl. Boxes show the median and interquartile range, whiskers the most extreme value within 1.5 × IQR, and points individual biological replicates. n = 6 biological replicates per group, from two independent experiments of three replicates each. Comparisons were made by exact two-sided Mann-Whitney U test, with p-values adjusted within each panel by the Benjamini-Hochberg procedure (* adjusted p < 0.05, ** adjusted p < 0.01; ns, not significant). In (F), four ΔdltA wells yielded no colonies and are plotted as zero; wild-type wells in this condition gave 1-6 colonies per spot, so counts were near the limit of detection.

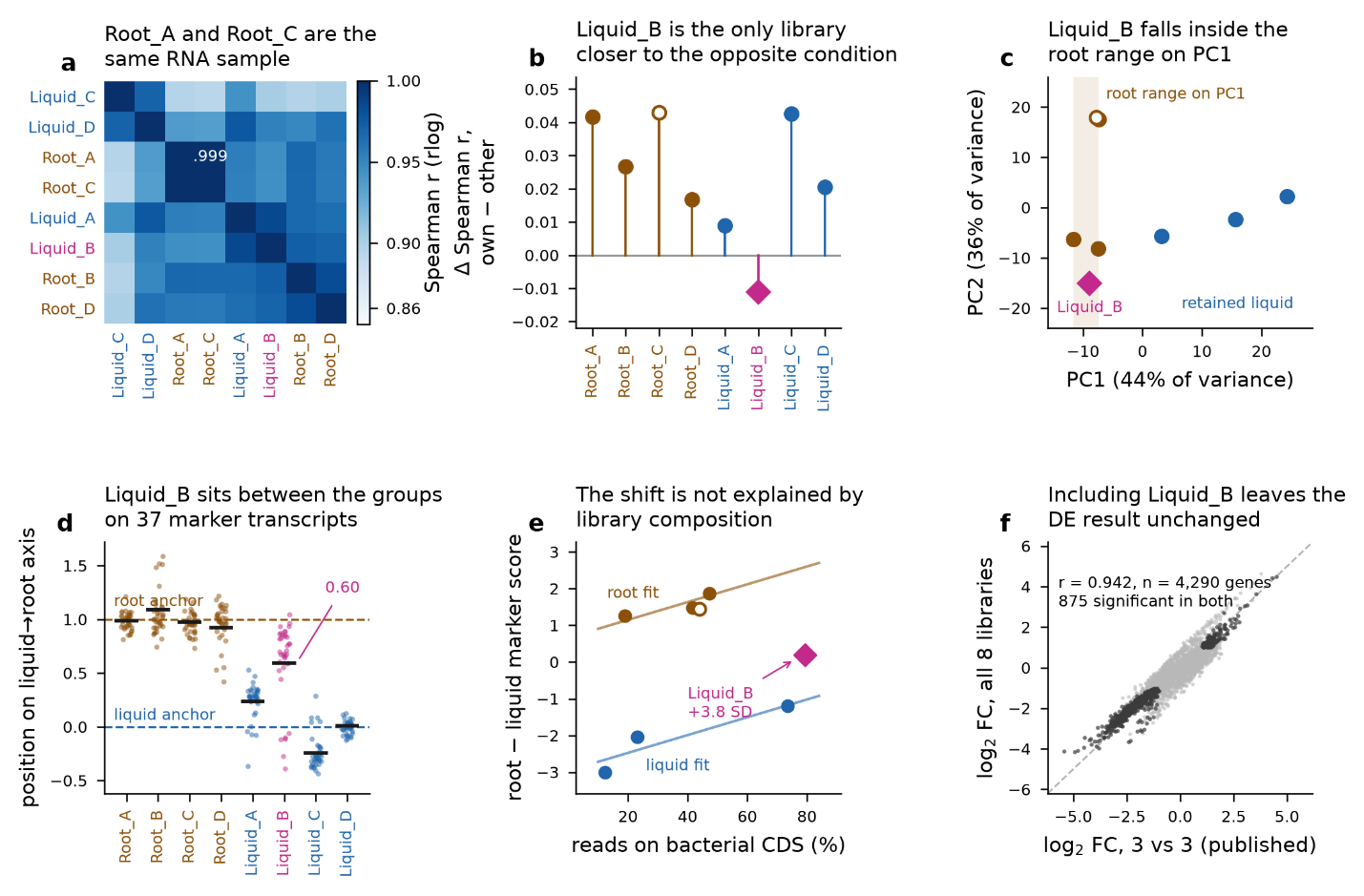

**Figure S11. Diagnostics for the exclusion of Root_C and Liquid_B from the root-versus-liquid RNA-seq comparison**. All panels use rlog-transformed counts for the 4,296 genes with non-zero total count across all eight libraries (DESeq2, blind dispersion estimation), except (F). Root libraries brown, liquid libraries blue; Root_C is drawn as an open circle and Liquid_B as a magenta diamond to mark them as excluded. **(A)** Spearman correlation between libraries, ordered by average-linkage clustering on 1 - r; Root_A and Root_C correlate at r = 0.999, identifying them as the same RNA sample. **(B)** For each library, mean correlation to the remaining members of its own condition group minus mean correlation to the opposite group; Liquid_B is the only library with a negative value. This statistic is sensitive to the composition of the comparison group and becomes positive (+0.011) if Liquid_C is omitted (Table Sxx). **(C)** Principal components of the 500 most variable genes; the shaded band spans the PC1 range of the four root libraries, which contains Liquid_B. **(D)** Position of each library on a liquid(0)–root(1) axis anchored on the six retained libraries, for the 37 signature genes with a root–liquid separation exceeding 1.5 rlog units (points, one per gene; black bar, whole-library score computed on all 100 signature genes). Dashed lines mark the anchor means. Liquid_B sits at 0.60, between the two groups, with a single unimodal distribution across markers. **(E)** Marker score against the fraction of reads assigned to bacterial coding sequence. Lines are a least-squares fit of score on condition plus coding-sequence fraction estimated from the six retained libraries only (n = 6, 3 residual degrees of freedom); Liquid_B lies 3.8 residual standard deviations above the liquid expectation at its own purity. **(F)** Log2 fold changes from the published three-versus-three contrast against a contrast including all eight libraries (n = 4,290 genes with an estimate in both; dark points, the 875 genes significant at padj < 0.05 in both; dashed line, identity).
